# Characterization of a novel type III CRISPR-Cas effector provides new insights into the allosteric activation and suppression of the Cas10 DNase

**DOI:** 10.1101/2019.12.17.879585

**Authors:** Jinzhong Lin, Mingxia Feng, Heping Zhang, Qunxin She

## Abstract

Antiviral defense by type III CRISPR-Cas systems relies on two distinct activities of their effectors: the RNA-activated DNA cleavage and synthesis of cyclic oligoadenylate. Both activities are featured as indiscriminate nucleic acid cleavage and subjected to the spatiotemporal regulation. To yield further insights into the involved mechanisms, we reconstituted LdCsm, a lactobacilli III-A system in *Escherichia coli*. Upon activation by target RNA, this immune system mediates robust DNA degradation but lacks the synthesis of cyclic oligoadenylates. Mutagenesis of the Csm3 and Cas10 conserved residues revealed that Csm3 and multiple structural domains in Cas10 function in the allosteric regulation to yield an active enzyme. Target RNAs carrying various truncations in the 3′ anti-tag were designed and tested for their influence on DNA binding and DNA cleavage of LdCsm. Three distinct ternary LdCsm complexes were identified. In particular, binding of target RNAs carrying a single nucleotide in the 3′ anti-tag to LdCsm yielded an active LdCsm DNase regardless whether the nucleotide shows a mismatch, as in the cognate target RNA (CTR), or a match in the noncognate target RNAs (NTR), to the 5’ tag of crRNA. In addition, further increasing the number of 3′ anti-tag in CTR facilitated the substrate binding and enhanced the substrate degradation whereas doing the same as in NTR gradually decreased the substrate binding and eventually shut off the DNA cleavage by the enzyme. Together, these results provide the mechanistic insights into the allosteric activation and repression of LdCsm enzymes.

## Introduction

CRISPR-Cas (clustered regularly interspaced short palindromic repeats, CRISPR-associated) systems constitute the adaptive and heritable immune system in bacteria and archaea, which mediates antiviral defense against invasive genetic elements in a small RNA-guided fashion^1–8^. The immune system consists of two parts: CRISPR arrays containing spacers derived from invading nucleic acids, and *cas* gene cassettes coding for enzymes or structural proteins that function in mediating the CRISPR immunity. CRISPR-Cas systems are classified into two broad classes based on the composition of their effector complexes: Those of Class 1 carry multi-subunit effectors and those of Class 2 possesses a single effector protein, and these antiviral systems are further divided into six main types (types I - VI) with >20 subtypes^9–11^.

Type III CRISPR systems are unique because they exhibit both RNA interference and DNA interference *in vivo* to protect their microbial hosts against invading nucleic acids^4,12–23^. Three activities are associated with these Type III immune systems, including target RNA cleavage^18,24–30^, target RNA-activated indiscriminate single-stranded (ss) DNA cleavage, a secondary DNase activity^15,19,20,31–37^ and synthesis of cyclic oligoadenylate (cOA), a second messenger that activates RNases of Csm6/Csx1 families, mediating cell dormancy or cell death^38–45^. Activation of the immunity requires mismatches between the 3′ anti-tag of a cognate target RNA (CTR) and the 5′ tag of crRNA whereas the full match between the 3′ anti-tag of non-cognate target RNA (NTR) and the 5′ tag of crRNA completely represses the immune response (see reviews^46–49^). Upon activation by the secondary messenger, Csx1/Csm6 RNases exhibit indiscriminate cleavage of viral and cellular RNAs, leading to cell dormancy or cell death to curb virus infection^38–43,50,51^. It is further believed that the type III CRISPR DNase eventually clears up remaining invading nucleic acids^52^, whereas the cOA secondary messenger is to be removed by ring nucleases^53^, allowing cells recover from the Type III immune response and restore the growth.

Structure of the *Streptococcus thermophilus* III-A (StCsm) effector complex has recently been resolved. This includes that of the StCsm binary complex and those of the StCsm^CTR^ and StCsm^NTR^ ternary effector complexes. The two ternary complexes have undergone major conformational change, relative to the binary one, but the active StCsm^CTR^ complex and the inactive StCsm^NTR^ complex show minimal conformational change except for the fact that the 3′ anti-tag of CTR and that of NTR are placed in different channels^54^. Very similar results were obtained from the structural analysis of the *Thermococcus onnurineus* Csm effector complex^55^ and the StCsm effector of different composition^56^. In particular, the active site of these Csm DNases exhibits little difference between the NTR-bound effector complex and the corresponding CTR-bound effector complex. As a result, the process of the activation of a binary Csm complex by CTR as well as its inhibition by NTR remains elusive.

To gain an further insight into the molecular mechanisms of the allosteric regulation of type III-A effector complexes, we characterized a Csm present in *Lactobacillus delbrueckii* subsp. *bulgaricus* (LdCsm). Its effector complex was reconstituted by expression in, and purified from, *Escherichia coli*. Investigation of the interactions between purified effector complexes and their target RNAs unravels an active ternary LdCsm complexes with CTR or NTR carrying a single nucleotide at their 3′ anti-tag regions, and these results provide a novel insight for allosteric activation and repression of the LdCsm DNase.

## Results

### *L. delbrueckii* subsp. *bulgaricus* encodes a novel III-A CRISPR-Cas system defective in cOA synthesis

The *L. delbrueckii* subsp. *bulgaricus* strain carries a type III-A CRISPR-Cas system (LdCsm) including a CRISPR array of 16 spacers (Fig. 1a), in addition to a type II CRISPR-Cas9 system in another chromosome location (GenBank: CP016393.1). This LdCsm system was chosen for characterization since phylogenetic analyses of a selected set of Cas10 proteins revealed that its Cas10, the *L. delbrueckii* subsp. *bulgarius* Csm1, is distantly related to the Cas10 proteins that have been studied thus far (Supplementary Fig. S1).

**Fig. 1.**
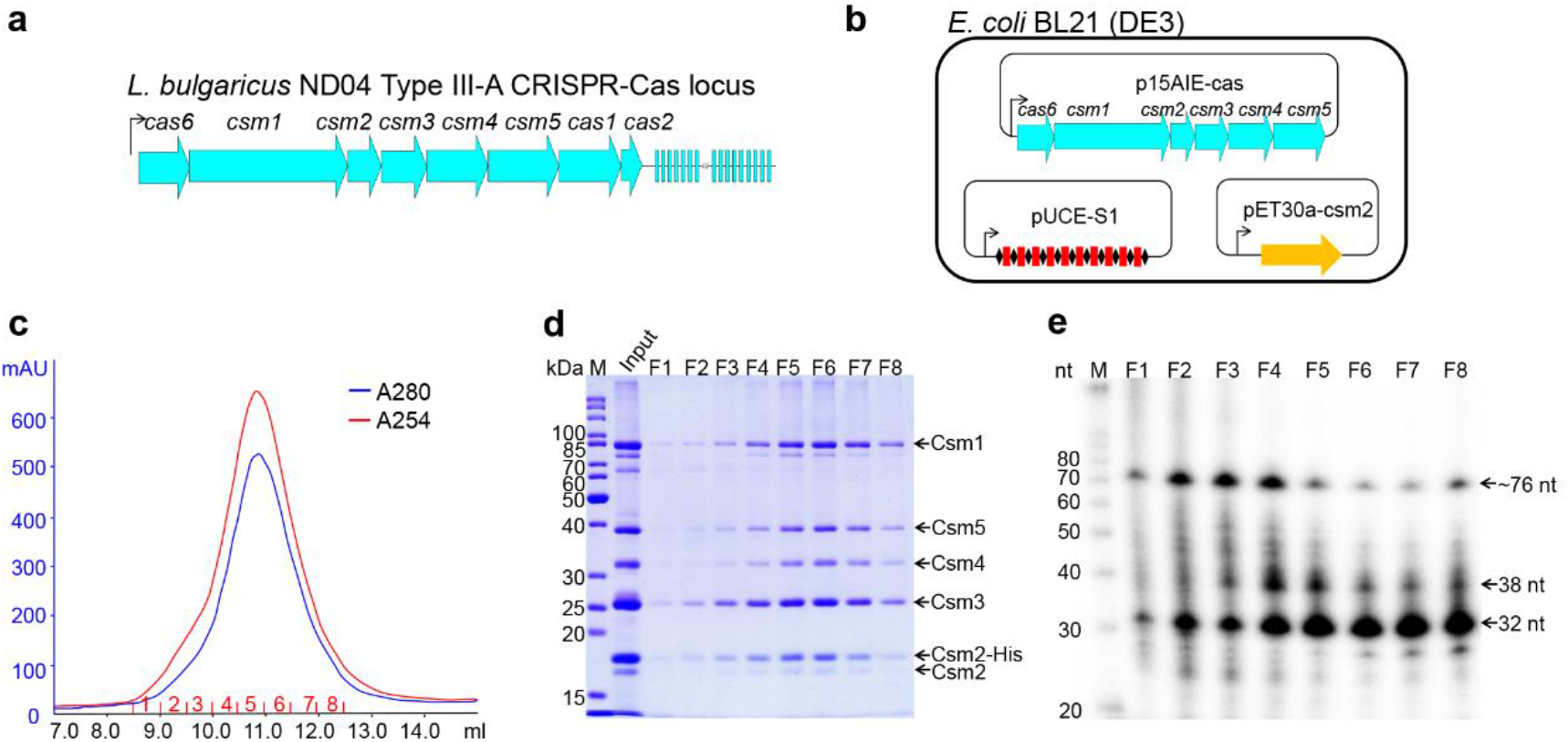
Cloning, expression and purification of the *L. delbrueckii* subsp. *bulgaricus* Csm complex in *E. coli*. (**a**) Schematic of the LdCsm system. *LdCsm* genes and the adjacent CRISPR assay are indicated with filled large arrows and small rectangles, respectively. Line with an arrowhead denotes the promoter of the *csm* gene cassette and the direction of transcription. (**b**) Strategy for reconstitution of the LdCsm effector in *E. coli*. *LdCsm* genes (*cas6*+*csm1-5* genes) were cloned into p15AIE, yielding p15AIE-cas (Supplementary Fig. S2). A CRISPR array carrying 10 copies of S1 spacer was generated and inserted into pUCE, giving pUCE-S1 (Supplementary Fig. S2). *LdCsm2* was cloned into pET30a, giving pET30a-Csm2 that yields the His-tagged Csm2 upon plasmid-born gene expression in the cell. All three plasmids were introduced into *E. coli* BL21(DE3) by electroporation. (**c**) UV spectrum of SEC purification of LdCsm effector complex. *E. coli* cell extracts were employed for Nickel-His tag affinity purification of LdCsm2 by which LdCsm effector complexes were copurified. The resulting protein samples were further purified by SEC. Blue: UV absorbance at 280 nm; red: UV absorbance at 254 nm. (**d**) SDS-PAGE analysis of SEC samples collected in the peak region. M: protein mass marker; Input: proteins purified by nickel Csm2-His affinity chromatography. (**e**) Denaturing gel electrophoresis of 5′-labeled RNAs from LdCsm samples. RNAs were extracted from the SEC-purified LdCsm samples. M: RNA size ladder.

We chose to reconstitute the LdCsm effector in *E. coli* since recombinant protein purification procedure has not been established for this lactobacillus yet. Three different *E. coli* vectors, i.e. p15AIE, pUCE and pET30a, were employed to clone all components of the immune system, including the complete set of III-A *cas* genes, a His-tag version of *csm2* and a synthetic CRISPR array (Supplementary Fig. S2). Expression of these plasmid-borne genes in the same cell yielded all this III-A Cas proteins, a His-tagged LdCsm2 protein and crRNAs (Fig. 1b), which were allowed to assemble into recombinant LdCsm ribonucleoprotein complexes in *E. coli*. The resulting effector complexes were then purified in two-step purification, the nickel-His tag affinity chromatography and the size exclusion chromatography (SEC). A single protein peak appeared in 10-12 ml in the SEC purification (Fig. 1c), indicative of copurification of large complexes. Analysis of these SEC samples by SDS-PAGE showed that each SEC sample contained 5 protein bands corresponding to the predicted sizes of Csm1, Csm2, Csm3, Csm4, and Csm5, respectively (Fig. 1d). The RNA component was extracted from the effector complexes using the Trizol agent. Denaturing PAGE analysis of extracted RNAs by radio-labelling and northern blot analysis of these RNAs by the radio-labeled DNA probe revealed 3 major RNA components of ~76, 38 and 32 nt. The largest species represented the complete unit of S1 crRNA carrying the 5′-repeat handle+S1 spacer+3′-repeat handle, representing the Cas6-cleaved crRNA product of a single spacer, whereas the two smaller RNA species were matured crRNAs (Fig. 1e and Supplementary Fig. S3). Together, these results indicated that LdCsm ribonucleoprotein complexes were reconstituted in *E. coli*. Since F5-F7 fractions mainly contained the LdCsm complex of 32 nt crRNA (Fig. 1e), the smallest ribonucleoprotein effector complex, they were pooled together, annotated as LdCsm and characterized.

To test if the *E. coli*-expressed effector could be active in RNA cleavage, LdCsm was mixed with four different radio-labeled RNAs individually, i.e. (a) a non-homologous RNA, S10, (b) the protospacer target RNA (PTR), S1-40, lacking any 3′ anti-tag, (c) the cognate target RNA (CTR), S1-46, exhibiting mismatches between its 6 nt 3′ anti-tag and the 5′ tag of the corresponding crRNA, and (d) the noncognate target RNA (NTR), S1-48, possessing the fully complementary sequence between its 3′ anti-tag and the 8 nt 5′ tag of the crRNA (Fig. 2a). These reaction mixtures were incubated at 37°C for 10 min and analyzed by denaturing PAGE. We found that, while the non-homologous RNA was not a substrate of LdCsm (Supplementary Fig. S4a), the effector cleaved all three homologous target RNAs in 6-nt periodicity with similar efficiencies. These results indicated that LdCsm possesses the backbone RNA cleavage activity and 3′ anti-tag sequences on target RNAs do not influence the RNA cleavage (Fig. 2b). Then, all three target RNAs were tested for their capability of mediating RNA-activated DNA degradation to S10-60, a single-stranded non-homologous DNA substrate (Supplementary Table S1). As shown in Fig. 2c, only CTR activated the LdCsm DNase, indicating that mismatches between the 3′ anti-tag of the target RNA and the corresponding 5′ tag of crRNA are essential for the activation. These results are in good agreement with those reported for other characterized type III CRISPR-Cas systems^15,17,19,20,31,34,35^. We further tested the minimal concentration of CTR is required for the ssDNase activity of LdCsm. To do that, a fluorophore quencher-labelled 16-nt poly-dT ssDNA reporter (FAM-poly-16T-BHQ1) containing a 5′-FAM (5′-Carboxyfluorescein) and 3′-BHQ1 (3′-Black Hole Quencher-1) was synthesized and employed as ssDNA substrate. Reactions containing 50 nM LdCsm and a wide range of concentrations of CTR (0 to 10 nM) were set up using a microplate. the result showed that 0.5~1 nM CTR is efficient to activate the ssDNase activity of LdCsm (Supplementary Fig. S5).

**Fig. 2.**
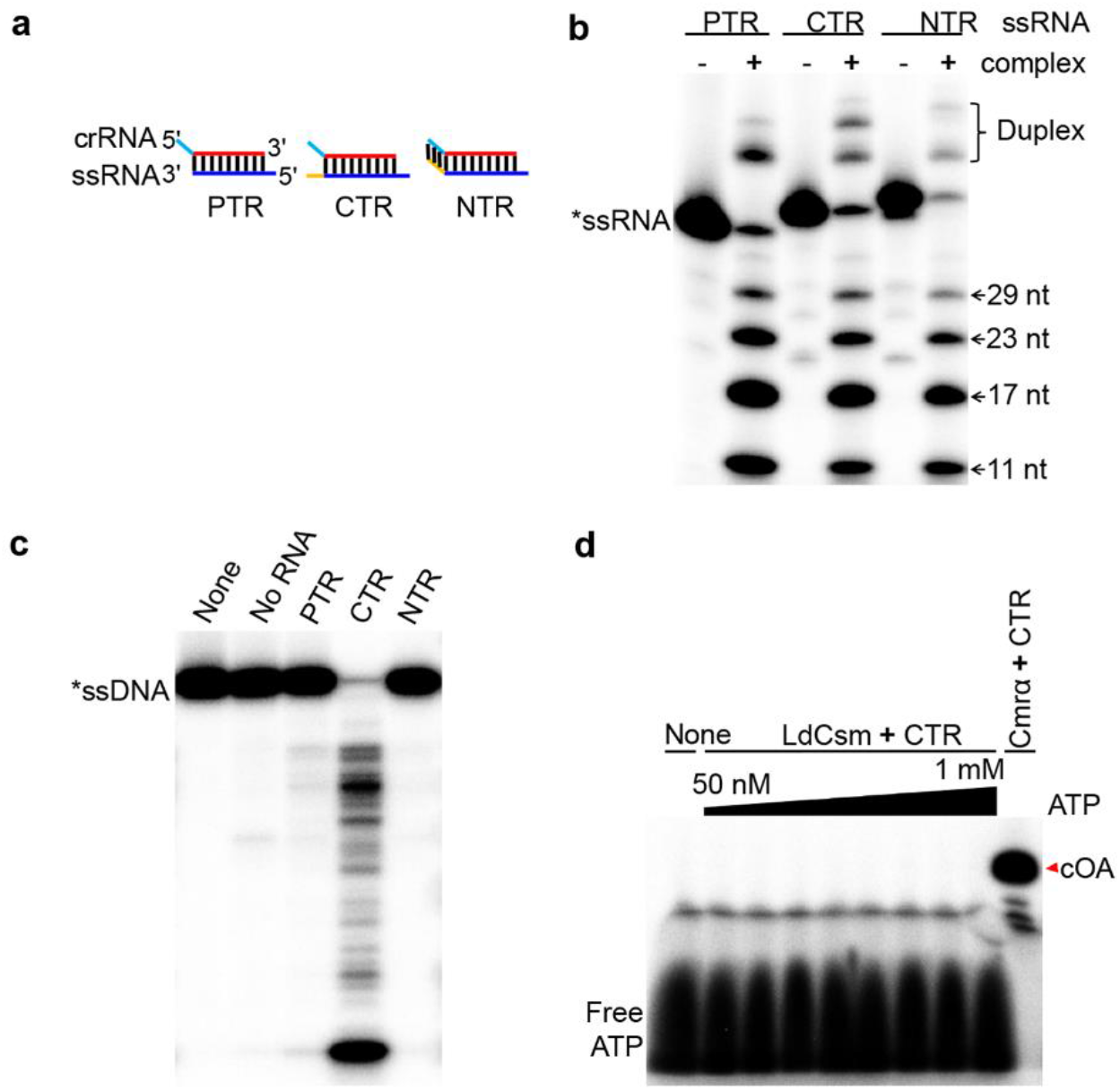
Biochemical characterization of the LdCsm effector complex. (**a**) Schematic of three different homologous target RNAs: CTR, cognate target RNA carrying 6-nt 3′ anti-tag with mismatch to the 5′ tag of the corresponding crRNA; NTR, noncognate target RNA containing 8-nt 3′ anti-tag that is complementary to the 5′ tag of the corresponding crRNA, and PTR, 40 nt protospacer target RNA completely lacking 3′ anti-tag. (**b**) Analysis of target RNA cleavage by LdCsm. Different target RNAs (50 nM) were individually mixed with 50 nM LdCsm and incubated for 10 min. The resulting samples were analyzed by denaturing PAGE. Duplex: Duplex of crRNA and substrate. (**c**) Analysis of RNA-activated ssDNA cleavage by LdCsm. 50 nM S10-60 ssDNA substrate was mixed with 50 nM LdCsm and 500 nM of each of the target RNA and incubated for 10 min. Samples were analyzed by denaturing PAGE. (**d**) Analysis of cOA synthesis by LdCsm. ~2 nM [α-^32^P]-ATP was mixed with a range of cold ATP (48 nM – 1 mM) and incubated with 50 nM LdCsm in the presence of 500 nM CTR for 120 min, the *S. islandicus* Cmr-α complex was used as the positive reference.

To date, all studied type III effectors are capable of producing cOAs, a secondary messenger that activates CRISPR-associated Rossmann fold (CARF) domain RNases of the Csm6/Csx1 family for general degradation of cellular RNAs^38–42^. To investigate if LdCsm could also do that, the effector was mixed with ATP and CTR in the presence of Mg^++^, and incubated for 2 h. Samples were analyzed by denaturing PAGE. As shown in Fig. 2d, whereas the *Sulfolobus islandicus* III-B Cmr-α (a positive reference^38^) consumed almost all 100 μM ATP for cOA synthesis, but LdCsm did not produce any detectable cOA in the same reaction setup. We repeated the experiments several times and also tested with a number of different metal ions, but constantly failed to detect any cOA in the ATP reaction (Supplementary Fig. S4f).

We noticed that LdCsm1 carries a <u>Q</u>GDD motif in Palm 2, the active site for cOA synthesis, which differs from the <u>G</u>GDD consensus (Supplementary Fig. S6a). To test if the occurrence of glutamine (Q597) residue in the motif could be responsible for cyclase inactivation in LdCsm, a Q597G substitution mutant was constructed to restore the common GGDD motif in LdCsm1, denoted Csm1^Q597G^. Characterization of the resultant mutated effector revealed that the GGDD-restored LdCsm still appeared to be inactive in cOA synthesis (Supplementary Fig. S4f). To this end, the LdCsm cyclase inactivation must not be resulted from any spontaneous mutation at the Cas10 cyclase active site.

### LdCsm system mediates anti-plasmid interference in *E. coli*

To investigate if the LdCsm system could mediate DNA interference *in vivo*, three test plasmids were constructed using pBad, an *E. coli* vector exhibiting arabinose-inducible expression. These included pBad-eGFP (pBad-G), a reference plasmid and two protospacer-carrying plasmids: one containing the CTR-S1 protospacer-GFP fusion gene (pBad-CTR), and the other possessing the NTR-S1 protospacer-eGFP (pBad-NTR) (Fig. 3a). Meanwhile, *E. coli* strain for the *in vivo* assay was generated by introduction of the expression plasmid p15AIE-Cas-S1 into *E. coli* BL21 (DE3) by electroporation. The rationale of the experiment is that plasmid-borne gene expression from p15AIE-Cas-S1 would yield binary LdCsm effector complexes in the host cells, which would mediate anti-plasmid activity if a cognate target RNA was to be expressed from a test plasmid. In addition, since the expression of CTR-S1 or NTR-S1 target RNA is controlled by the Bad promoter that confers very stringent arabinose-inducible expression, target RNAs would only be synthesized in L-arabinose media but not in glucose media. Therefore, examination of colony formation efficiency of each plasmid on both arabinose plates and glucose media would reveal *in vivo* transcription-dependent anti-plasmid interference of the immune systems to be tested (Fig. 3b).

**Fig. 3.**
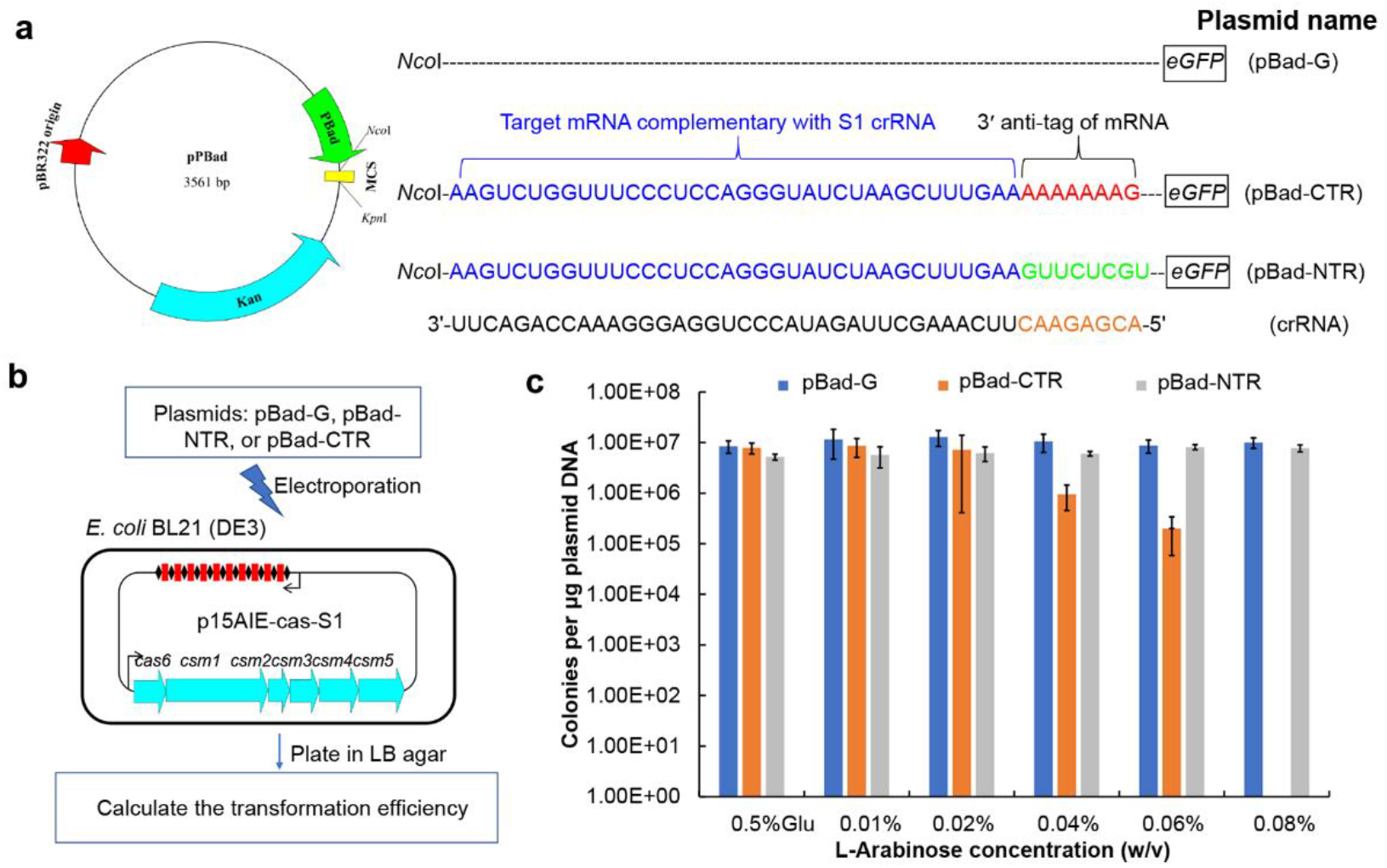
Anti-plasmid activity of the LdCsm system. (**a**) Schematic of the test plasmids pBad-G, pBad-CTR and pBad-NTR. Gene expression from these plasmids is under the control of a L-arabinose-inducible Bad promoter. Therefore, mRNAs carrying CTR or NTR are expressed from the corresponding plasmids (pBad-CTR or pBad-NTR) in the presence of L-arabinose but their expression is completely repressed in glucose media. (**b**) Schematic of the interference plasmid assay for determination of the LdCsm anti-plasmid activity. (**c**) Transformation efficiency data of the three different plasmids obtained from different growth media.

All test plasmids were introduced into the genetic host by transformation, and transformed *E. coli* cells were plated on 6 different medium plates containing 0.5% glucose or arabinose of different contents (0.01-0.08%). Following transformation data were obtained: (a) pBad-G and pBad-NTR produced very similar numbers of transformants on all 6 growth media, (b) pBad-CTR gave very similar transformation efficiency data on plates containing glucose or very low concentrations of L-arabinose (0.01 or 0.02%), (c) increasing the L-arabinose content in the medium to 0.04% and 0.06%, yielded ca. 10 and 100-fold decrease in transformation efficiency, and (d) the presence of 0.08% inducer completely abolished colony formation by pBad-CTR-containing cells (Fig. 3c). These results indicated, while NTR can effectively turn off the LdCsm immunity, CTR is capable of conferring plasmid clearance to the LdCsm system albeit it only possesses the RNA-activated ssDNA cleavage activity. This is in contrast to the current knowledge of antiviral mechanisms by type III CRISPR-Cas systems in which the cOA signaling pathway plays a main role in the immunity^13,51,52,57,58^, suggesting LdCsm represents the novel type III CRISPR-Cas system, which exhibits the robust RNA activated DNase activity to resist the invasive plasmid.

### Multiple Cas10 domains contribute to the LdCsm DNase activity

The largest subunit Cas10 in type III-A effector complexes contains 4 conserved domains, including a HD domain, two Palm domains (Palm1 and Palm2), a Linker and a D4 structural region (Fig. 4a). Biochemical and structural analyses of different type III CRISPR-Cas systems have revealed that the HD domain is responsible for ssDNA cleavage^15,19,20,31,34,35^, Palm1 and Palm2 domains function in cOA synthesis^38–41,43^ whereas the Linker domain plays a regulatory role in both target RNA-activated DNA cleavage and cOA synthesis^54^. However, it remains unclear whether a concerted action of different Cas10 domains would be required to yield Csm DNase with optimal activity. The robustness of the LdCsm DNase rendered it a good system for this characterization.

**Fig. 4.**
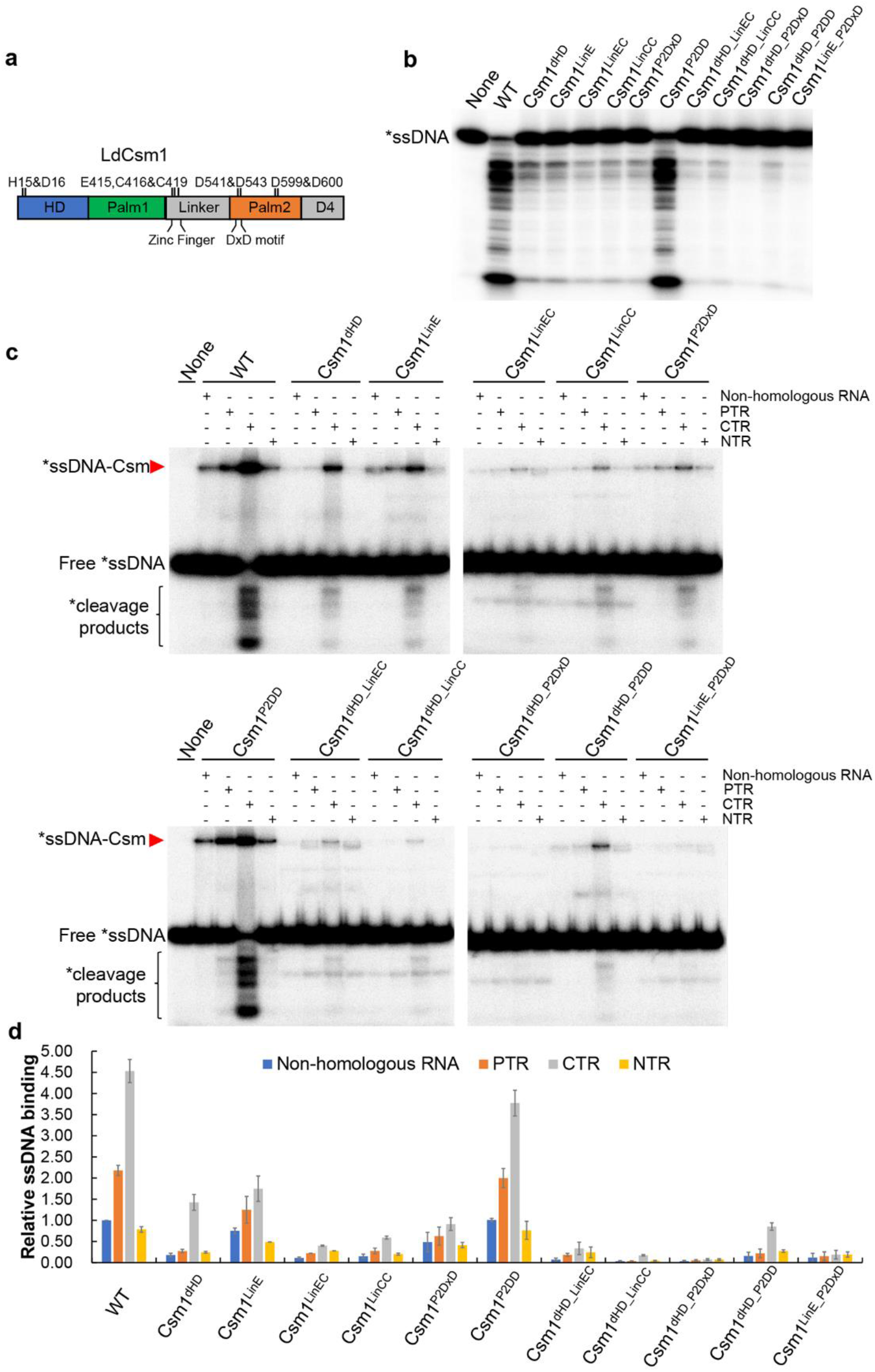
Effect of LdCsm1 mutations on ssDNA binding and cleavage by the LdCsm effector complex. (**a**) Domain architecture of the LdCsm1 protein. HD represents the HD-type nuclease domain; Palm 1 and Palm 2 denote the two cyclase domains; Linker is a domain that adjoins the Palm1 and Palm2 domains, consisting 4 cysteine residues, D4 is located in the C-terminus rich in α-helices. Amino acid residues selected for alanine substitution mutagenesis are indicated with their names and positions. (**b**) RNA-activated ssDNA cleavage by effectors carrying one of the constructed LdCsm1 mutants. 50 nM S10-60 ssDNA substrates were mixed with 50 nM mutated LdCsm carrying each of LdCsm1 mutant proteins and 500 nM CTR, and incubated for 10 min. Samples were analyzed by denaturing PAGE. (**c**) ssDNA binding by effectors carrying each of the constructed LdCsm1 mutants. 5 nM labeled S10-60 ssDNA were incubated with 100 nM of LdCsm effectors indicated in each experiment. 400 nM of non-homologous RNA (S10 RNA) or in the presence of 500 nM of one of the target RNAs, PTR or CTR or NTR for 3 min. Samples were analyzed by non-denaturing PAGE. Red arrowheads indicate the Csm-ssDNA complex. (**d**) Relative ssDNA binding between the wild-type LdCsm effector and its LdCsm1 mutated derivatives. The relative ssDNA binding activities were estimated by image quantification of the non-denaturing PAGE in (c) by the accessory analysis tool in Typhoon FLA 7000, the ssDNA activity of LdCsm in non-homologous RNA was used as the standard and set up as 1. Results shown are average of three independent assays, bars represent the mean standard deviation (± SD).

Nine conserved amino acids present in the HD, Linker, and Palm2 domains of LdCsm1 were chosen for substitution mutagenesis, giving 6 single domain mutants. These included Csm1^dHD^, a HD domain mutant carrying H15A and D16A double substitution, 3 Linker domain mutants, i.e. Csm1^LinE^ (E415A substitution), Csm1^LinEC^ (E415A and C416A substitutions), and Csm1^LinCC^ (C416A and C419A mutations), and 2 Palm2 domain mutants, Csm1^P2DxD^ (D541A and D543A substitutions) and Csm1^P2DD^ (D599A and D600A mutations). Then, some mutated motifs were combined, yielding double domain mutations including two HD/Linker mutants (Csm1^dHD_LinEC^ and Csm1^dHD_LinCC^), two HD/Palm2 mutants (Csm1^dHD_P2DxD^ and Csm1^dHD_P2DD^) and Csm1^LinE_P2DxD^, a Linker and a Palm2 mutant (Supplementary Table S2).

Each mutated *csm1* gene was used to replace the wild-type *csm1* on p15AIE-cas, and the resulting plasmids were introduced into *E. coli* to express LdCsm effectors carrying each of the mutated LdCsm1 protein. These mutated LdCsm effector complexes were obtained as described for the wild-type LdCsm effector. The protein components of these effector complexes were checked by SDS-PAGE, and this showed that they all contained 5 different Csm subunits as for the wild-type LdCsm complex (Supplementary Fig. S6b), indicating that none of these Csm1 mutations affected the effector assembly. These effector complexes were then tested for target RNA cleavage using S1-46, the CTR, as the substrate and they all exhibited the backbone cleavage (Supplementary Fig. S6c).

Next, RNA-activated DNase activity was examined for each mutated complex in reactions containing 50 nM effector, 50 nM ssDNA and 500 nM CTR. For single domain mutants, we found that all mutations (except for Csm1^P2DD^) strongly impaired the ssDNase activity, including Csm1^dHD^, Csm1^LinE^, Csm1^LinEC^ Csm1^LinCC^ and Csm1^P2DxD^ (Fig. 4b). These results indicated that, in additional to the HD domain, the putative cleavage site, three other Csm1 motifs, i.e. E415 and C416 C419 in the Linker region and DxD in the Palm2 domain are also important for the LdCsm DNase. Analysis of the double motif mutants further showed, while combined mutations of Csm1^dHD_LinEC^, Csm1^dHD_LinCC^ and Csm1^dHD_P2DD^ possessed similar DNase activities relative to those of single motif mutants, those of Csm1^dHD_P2DxD^ and Csm1^LinE_P2DxD^ completely abolished the LdCsm DNase cleavage (Fig. 4b). To this end, we concluded that three Cas10 motifs, including HD, E415, C416 and C419 (Zinc finger) in the Linker as well as D451and D453 (DxD) in Plam2, play critical roles in the LdCsm DNA cleavage.

To yield a further insight into the LdCsm DNA cleavage, all these Csm1-mutated effector complexes were tested for substrate binding in which 25, 50 or 100 nM effector complex was mixed with 500 nM CTR and 5 nM radio-labeled S1-60 ssDNA. After incubation for 3 min, the formation of LdCsm-ssDNA complexes was checked by nondenaturing PAGE. As shown in Supplementary Fig. S6e, the substrate affinity of these effector complexes fell into three different categories. First, wild-type (WT) and LdCsm_Csm1^P2DD^ showed a strong binding (100% and ~80%), which is consistent with their unimpaired DNase activity; second, LdCsm_Csm1^dHD^ and LdCsm_Csm1^P2DxD^ retained ~30% of the binding capacity of the wild-type LdCsm effector, whereas the last group included those with mutations in Linker, only exhibiting 5-20% of substrate binding. Taken together, these data indicated that the HD motif, the Zinc finger and the DxD motif function in facilitating substrate binding of the LdCsm effector complex.

### LdCsm1 Linker domain and Palm2 DxD motif function in the allosteric control of the LdCsm DNase

The establishment of ssDNA binding assay with LdCsm prompted us to investigate how target RNAs could regulate the LdCsm DNase. For this purpose, 400 nM, S10 RNA (non-homologous RNA) and 500 nM of PTR, NTR or CTR was mixed with 100 nM LdCsm and 5 nM of labeled S1-60 DNA individually. After incubation at 37 °C for 3 min, samples were analyzed by non-denaturing PAGE. We found that, while the WT LdCsm effector showed little DNA-binding activity in the presence of non-homologous RNA or NTR, PTR greatly enhanced the DNA binding, and that activity was further facilitated by CTR for ca. 4-fold (Fig. 4c & 4d), indicating that CTR induces allosteric regulation on substrate binding of LdCsm. These results also indicated that the activity of LdCsm DNase involves the protospacer binding-induced allosteric regulation, in which protospacer region-bound facilitates the ssDNA substrate binding but the complexes remain inactive, and CTR-dependent activation of the LdCsm DNase. Therefore, 3′ anti-tag of both CTR and NTR regulate LdCsm DNA binding.

Analyses of all above Csm1-mutated effector complexes revealed that, while five mutated effectors (LdCsm_Csm1^P2DD^, LdCsm_Csm1^dHD^, LdCsm_Csm1^LinCC^, LdCsm_Csm1^dHD_LinCC^, and LdCsm_Csm1^dHD_P2DD^) exhibited a pattern of target RNA activation that is very similar to that of the wild-type LdCsm, those carrying E415A and/or alanine substitutions in Palm2 DxD did not show such a significant CTR-enhanced ssDNA substrate binding; their CTR and PTR ternary complexes basically remained inactive (Fig. 4c & 4d). These results indicated that among the three motifs essential for the LdCsm DNase, HD is not involved in CTR-induced allosteric regulation of LdCsm whereas both E415 and P2-DxD are essential for the allosteric regulation.

Taken together, our results unraveled two distinctive functions for the Cas10 protein: (a) consistent with the results obtained with other type III effectors, the HD motif of LdCsm1 hosts the catalytic site of the DNase; (b) E415 and P2-DxD are involved in mediating allosteric regulation of the ternary effector to yield active enzyme.

### Identification of Csm3 amino acids involved in the regulation of the LdCsm DNase

The ssDNA substrate binding assay showed that PTR-bound ternary LdCsm showed a higher substrate binding compared with that in the presence of non-homologous RNA (Fig. 4c), indicating that the protospacer region-bound drives certain regulation of LdCsm complex. To yield an further insight into the mechanism, some Csm3 variants were constructed and the mutated effector complexes were purified and characterized *in vitro*.

Alignment of a selected set of Csm3 proteins revealed a number of conserved amino acids among which may have a structural function or form the intermolecular contacts with crRNA or target RNA (Supplementary Fig. S7). There are three conserved residues: H20, D34 and D106 do not play a role in structure or crRNA or target RNA binding (LdCsm3 number). D34 was the predicted active site of the LdCsm RNase whereas functions of the remaining two amino acids were uncertain. To study that, we constructed several LdCsm3 mutants, including alanine substitution of H20, D34, or D106 as well as double substitutions (H20/D34 and D34/D106). LdCsm effector complexes carrying each of these Csm3 mutations (designated Csm3^H20A^, Csm3^D34A^, Csm3^D106A^, Csm3^H20/D34-A^ and Csm3^D34/D106-A^) were purified from *E. coli* and characterized. Target RNA cleavage assay showed that, while neither Csm3^H20A^ nor Csm3^D106A^ substitutions influenced the target RNA cleavage, Csm3^D34A^ substitution greatly impaired the RNA cleavage of the effector (Fig. 5a). These results confirmed that the D34 is the active site of target RNA cleavage of the LdCsm system as demonstrated for the conserved aspartic acid residue in all other known type III CRISPR-Cas systems. Analysis of RNA-activated ssDNA cleavage showed that, the LdCsm complex containing the Csm3^D34A^ substitution exhibited higher ssDNase activity relative to WT complex, DNase activity was strongly impaired in the remaining 4 mutated Csm3 complexes (Fig. 5b).

**Fig. 5.**
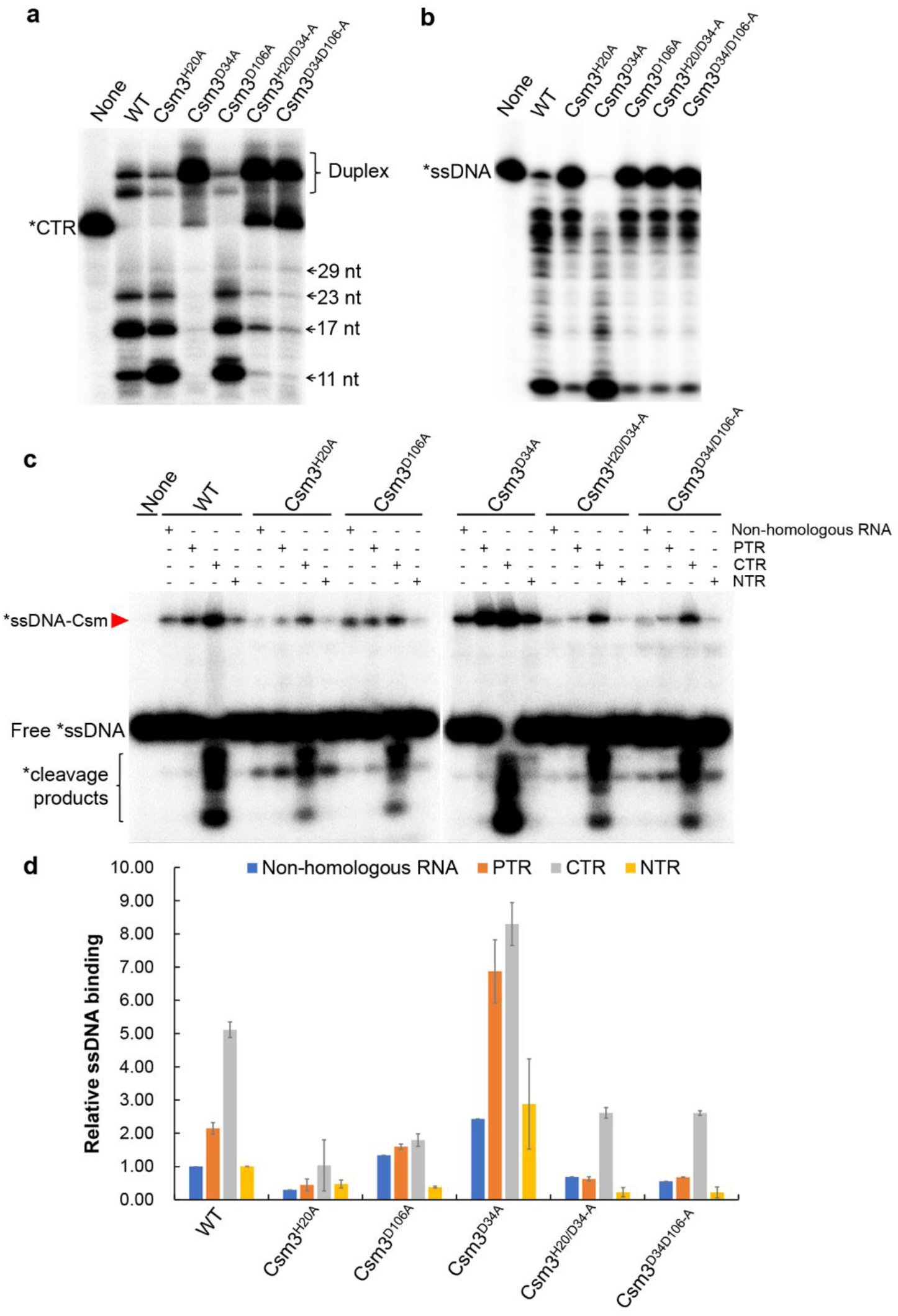
Effect of LdCsm3 mutations on the ssDNA cleavage and binding of LdCsm. (**a**) Target RNA cleavage of LdCsm3 mutated derivatives. 50 nM of S1-46 RNA were incubated with 50 nM of LdCsm or the indicated mutant derivatives for 10 min and the samples were analyzed by denaturing PAGE. Duplex: Duplex of crRNA and substrate. (**b**) RNA-activated ssDNA cleavage by effectors carrying one of the constructed LdCsm3 mutants. Reaction conditions were the same as in Fig. 4b. (**c**) ssDNA binding by effectors carrying each of the constructed LdCsm3 mutants. Reaction conditions were the same as in Fig. 4c. (**d**) Relative ssDNA binding between the wild-type LdCsm effector and its LdCsm3 mutated derivatives. The ssDNA activity of LdCsm in non-homologous RNA was used as the standard and set up as 1. Results shown are average of three independent assays, bars represent the mean standard deviation (± SD).

To test their spatiotemporal regulation, these Csm3 mutated complexes were incubated with the ssDNA substrate in the presence of different RNAs. We found that, except for Csm3^H20A^ substitution exhibiting an impaired ssDNA binding for all tested RNAs, Csm3^D34A^ exhibited a higher ssDNA substrate binding enhancement in presence of PTR vs. non-homologous RNA than WT complex (ca. 3-fold vs. 2-fold, respectively), which might due to the impaired target RNA cleavage allows a longer persistence of the ternary complex. However, Csm3^D106A^ substitution and the double substitutions Csm3^H20/D34-A^ and Csm3^D34/D106-A^ did not show such a significant PTR-induced ssDNA substrate binding enhancement relative to WT complex (Fig. 5c & 5d). Together, These results confirmed that CTR-activated LdCsm ssDNase activity involves the protospacer binding-induced allosteric regulation, consistent with the results obtained from structural analysis with other Csm or Cmr complexes^54,55,59^. These results also indicated that two Csm3 residues, H20 and D106, are involved in mediating regulation of the ternary LdCsm complex to yield active enzyme.

### Identification of different target RNA-LdCsm ternary complexes with distinctive substrate binding and DNA cleavage

To investigate how target RNA could activate the LdCsm DNase, 3’-truncation variants of the cognate target RNA were generated, carrying +1, +2, +3, +4, +5, +6 nt of the 3′ anti-tag region of CTR (annotated as CTR^+1^ to CTR^+6^) as well as the full length CTR (CTR^Full^) (Supplementary Table S1). Each target RNA was tested for its capability to facilitate DNA binding and DNA cleavage by LdCsm. As shown in Fig. 6a, while the binary LdCsm showed little DNA binding, PTR, the target RNA lacking any 3′ anti-tag sequence, greatly facilitated the substrate binding of the LdCsm complex (ca. 55% of CTR-LdCsm) but it failed to mediate DNA cleavage. Next, when CTR1, a target RNA that extended the PTR by a nucleotide at the 3′ anti-tag position, formed a ternary effector complex with LdCsm, the resulting effector not only showed further increased substrate binding activity (ca. 90% of the full activity) but also activated for the DNA cleavage (ca. 35% of full activity). This suggested that a single nucleotide at the 3′ anti-tag of CTR is efficient to convert the inactive ternary LdCsm complex to an active one. Finally, DNA cleavage activity by the LdCsm effector peaked with CTR^+4^ that carries 4 nt 3′ anti-tag of CTR, which could have completed the allosteric regulation of the immune system (Fig. 6a)

**Fig. 6.**
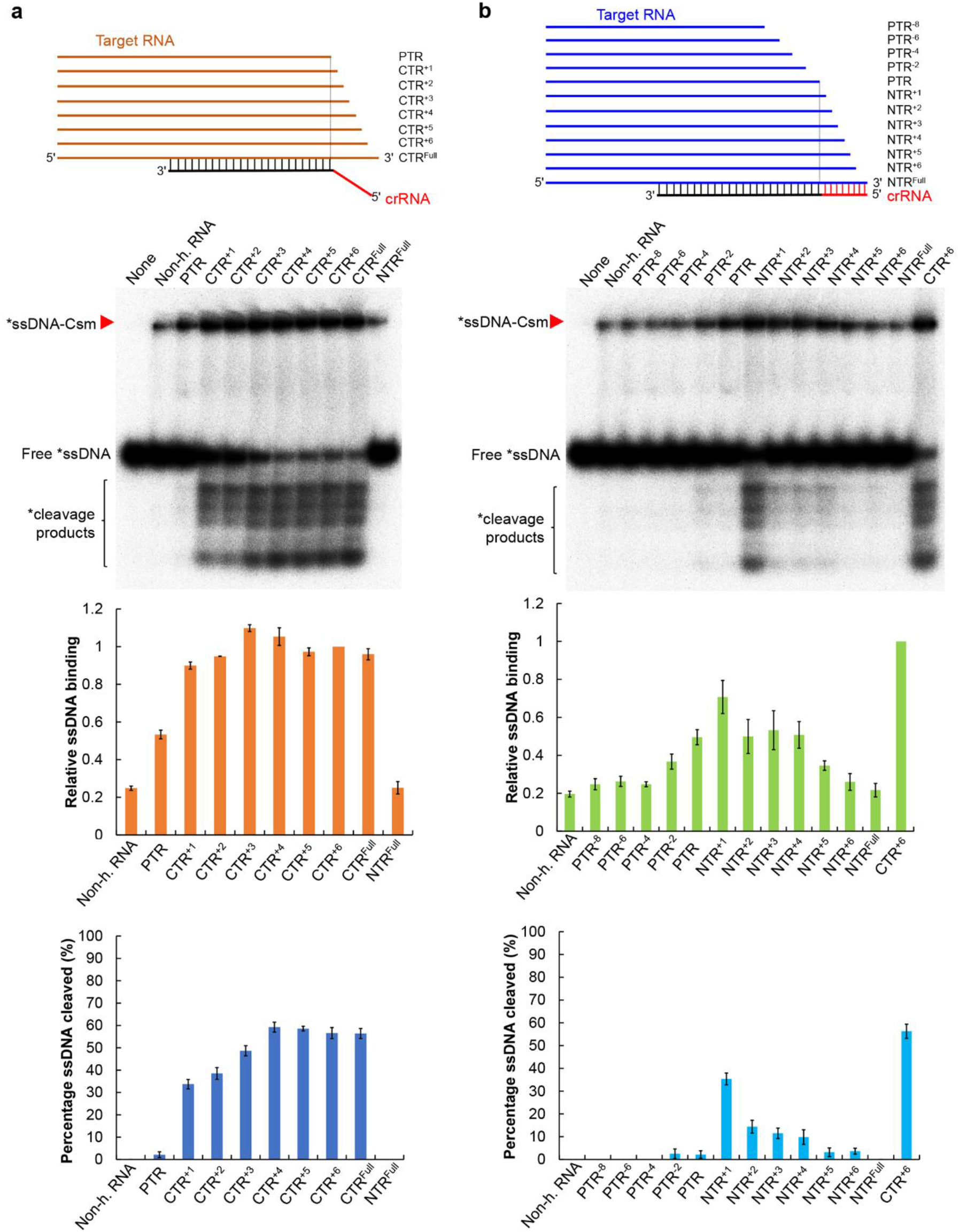
Target RNA-directed allosteric regulation of LdCsm involves activation and deactivation mechanisms. (**a**) CTR activates the LdCsm DNase. (**b**) NTR mediates autoimmunity avoidance by deactivation. Reactions were set up with 5 nM S10-60 ssDNA, 100 nM of LdCsm and 400 nM non-homologous RNA (non-h. RNA) or 500 nM target RNA. After addition of one of target RNAs, the mixture was incubated at 37 °C for 3 min. Samples were then analyzed by non-denaturing PAGE (in this page) or denaturing PAGE (Supplementary Fig. S8). Red arrowheads indicate the LdCsm-ssDNA complex. Relative ssDNA binding and percentage ssDNA cleaved of LdCsm facilitated by each of these target RNAs were estimated by image quantification of bands on non-denaturing PAGE and denaturing PAGE, using the accessory analysis tool equipped with a Typhoon FLA 7000. For the quantification of the substrate binding, the amount of ssDNA-LdCsm-CTR^+6^ complex is arbitrarily defined as 1. Results of average of three independent assays are shown with bars representing the standard deviation (± SD).

### NTR inhibits the LdCsm DNase by preventing substrate binding

The same approach was then employed to investigate how NTR could inhibit the LdCsm DNase. Truncated derivatives of NTR were generated (i.e. NTR^+1^ through NTR^+6^) as for the CTR derivatives, and in addition, a few PTR truncations (PTR^−8^ to PTR^−2^) were also made (Supplementary Table S1). These target RNAs were tested for their capability of facilitating substrate binding and cleavage as above described. We found that PTR and its truncated derivatives (PTR^−4^ to PTR^−8^) showed marginal differences both in ssDNA binding and in ssDNA cleavage, suggesting that these ternary complexes could belong to the same category, equivalent to the stage of the binary LdCsm effector (Fig. 6b). While PTR^−2^ and PTR showed similar effects on ssDNA binding, NTR^+1^, however, mediated a major stimulation to the LdCsm DNA binding (ca. 70% of the full activity), and it also facilitated the DNA cleavage (ca. 35% of full activity), and in fact, the stimulating effect by NTR^+1^ is comparable to that by CTR^+1^ (Fig. 6). These results not only implied the first nucleotide at the 3′ anti-tag of NTR can still allow activation of the HD nuclease domain, but also indicated that the process of repression by NTR shares some common activation with the CTR-dependent activation process of LdCsm DNase. Thereafter, extension of the 3′ anti-tag of NTR greatly reduces the DNA binding and gradually decreases DNA cleavage (Fig. 6b), suggesting that the interaction between 3′ anti-tag of NTR and the 5′ tag of crRNA could have deactivated the enzyme by restricting the accessibility of the ssDNA substrate to the active site.

## Discussion

Type III CRISPR-Cas systems characterized thus far show three different interference activities: target RNA cleavage, RNA-activated indiscriminate DNA cleavage and cOA synthesis among which the latter two activities are responsible for the DNA interference by these systems whereas target RNA cleavage plays a regulatory role (see reviews^46,47,49^). Here we report a type III-A subtype CRISPR-Cas system that shows robust RNA-activated ssDNA cleavage, but lacking in cOA synthesis. This is consistent with the fact that the genome of the *L. delbrueckii* subsp. *bulgaricus* genome does not code for any detectable Csm6 homologs. Interestingly, we find that a plasmid-borne LdCsm system is sufficient to mediate interference plasmid clearance in *E. coli*. This suggests that the LdCsm system probably only utilizes the RNA-activated ssDNase to mediate the antiviral immunity in the original host. We have also explored the simplicity of the LdCsm antiviral mechanism for investigating mechanisms of the DNA cleavage by type III CRISPR-Cas systems and our research has yielded mechanistic insights into the target RNA-mediated activation and repression of the LdCsm DNase.

Structures of several type III effector complexes have been solved^54–56,59,60^ and these analyses have revealed that formation of target RNA-Csm ternary effectors involves a minimal conformational change that is very similar between the CTR ternary effectors and the corresponding NTR ternary ones although only the former has been activated for DNA cleavage^54–56^. As a result, there is no major structural difference at the catalytic site of active versus inactive Csm DNases, and the only difference is the 3′ anti-tag sequences of the two different types of target RNAs are individually placed in different channels in their structures^54^. Furthermore, in the structure of the PTR-StCsm complex, major conformational change also occurs although the target RNA lacking any 3′ anti-tag. Since structure of a DNA substrate-bound Csm effector complex has not been resolved, how these effectors interact with their DNA substrates remains to unknown. For this reason, the only biochemical criterion to distinguish different effector complexes is to analyze their DNA cleavage activity. In this work, we have established a ssDNA binding assay for LdCsm, and our detailed analysis of substrate binding for this immune system has revealed that different target RNAs are capable of facilitating differential substrate binding and DNA cleavage to the LdCsm effector. Thus, it is very interesting to study how allosteric regulation differentially influences the two activities of the LdCsm DNase.

Structural analysis of StCsm by You et al.^54^ has indicated that the Cas10 Linker domain functions in mediating conformational change. Here, we find that that both the DxD motif in Palm 2 and the E415 residue in the Linker domain play an essential role in mediating the allosteric regulation of the LdCsm complex *in vitro*. To date, detailed functions of the Cas10 linker E415 and P2 DxD motifs have not been studied for other type III CRISPR-Cas systems although all three amino acids are well conserved in Cas10 proteins (Supplementary Fig. S6a). Therefore, it is of a great interest to investigate if their functions revealed for LdCsm represent a general mechanism for all type III immune systems.

In this paper, we have presented the detailed biochemical analysis of substrate binding by LdCsm effector complex, and our analysis shows that the Cas10 Zinc finger modulates the enzyme substrate binding since E415A substitution yielded an enzyme with greatly reduced DNA binding and inactive in DNA cleavage (Fig. 4). As E415 is probably part of the Zinc finger motif, this suggests that the Cas10 Zinc finger domain can have dual function in this immune system, namely both as an executor to the allosteric regulation and as a regulator to substrate binding^46,61^. The other study on substrate binding was conducted by single-molecule fluorescence microscopy analysis of *Staphylococcus epidermidis* Csm by Wang et al^62^. They find that Cas10 subunit is locked in a static configuration upon NTR binding in which the DNA binding pocket of the effector appears to be in a closed form, inaccessible to substrate. However, upon CTR binding, Cas10 exhibits a larger conformational space in the active site^62^. Together with our biochemical data, it is plausible to predict that there exists a substrate-binding pocket in the DNase of type III effector complexes although this structural domain remains to be illustrated by their structural analysis.

The possibility of characterization of effector complexes for both substrate binding and DNA cleavage allows us to identify target RNA-LdCsm effector complexes exhibiting differential activities, and we propose that these effector complexes probably represent intermediates for structural analysis to study the molecular mechanisms of the CTR activation or NTR repression of the LdCsm system (Fig. 7). These include: (a) The LdCsm-PTR complex showing an elevated level of substrate binding but inactive in catalysis, (b) the LdCsm-CTR^+1^ and LdCsm-NTR^+1^ exhibiting most of substrate binding and catalysis activities, representing another type of intermediates. There are two apparent features with the two effectors: they differ from LdCsm-PTR only by the first nucleotide present in the 3′ anti-tag of target RNAs, and the nucleotide in CTR is different from that in NTR. Thus, this nucleotide could functions as a trigger to induce the major allostery in LdCsm ternary to yield an active effector complex and the interaction between the effector with the nucleotide should not be specific, (c) LdCsm-CTR^+4^ represents another effector complex that should have adopted the same conformation since it exhibits the maximal substrate binding and catalysis, and finally (d) and LdCsm-NTR^Full^ represents the inactive conformation of the effector completely deactivate the LdCsm complex, ensuring autoimmunity avoidance of the LdCsm system (Fig. 7). These LdCsm effector complexes provide good model for structural analysis to reveal the molecular mechanisms of activation and repression of Csm DNases.

**Fig. 7.**
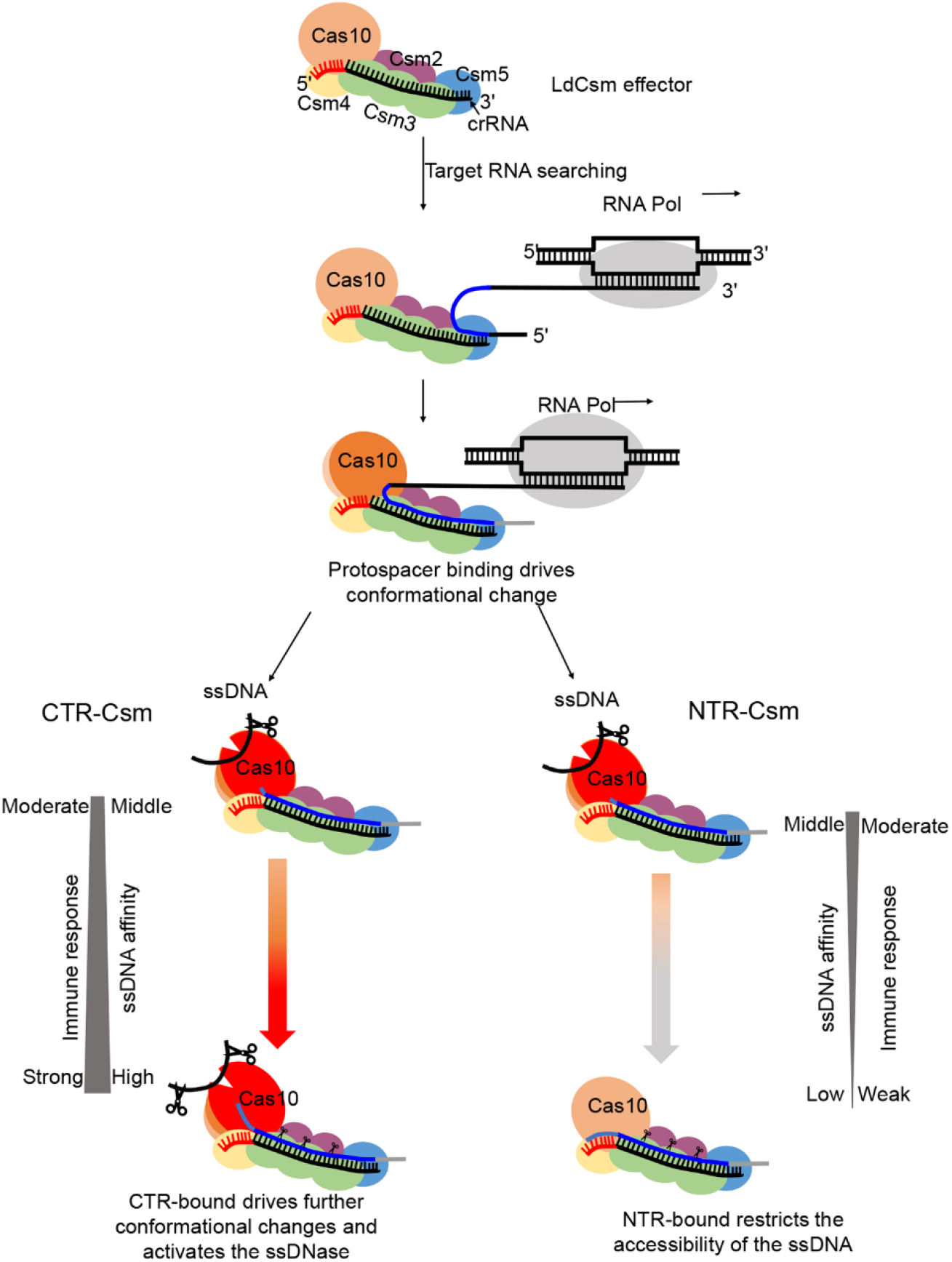
Model of allosteric activation and repression of the LdCsm DNase. The previous works have proposed that the initial recognition of nascent transcript at the 5′ end of target RNA for type III complex, since both of Csm5 subunit in Csm complex and Cmr1 subunit in Cmr complex are crucial for target RNA binding^36,37,68^. These suggested that the binary LdCsm effector complex interacts with target transcript initially at the 5′ end of target RNA and further via sequence complementarity between the protospacer and the corresponding crRNA, leading to the formation of a ternary effector complex with a major conformational change. Addition of a single nucleotide at the 3′-end of protospacer RNA results in an important allosteric change in the LdCsm DNase, giving an active enzyme. CTR-bound LdCsm exhibits the full level of substrate binding and DNA cleavage whereas NTR-bound LdCsm closes the substrate-binding pocket, which deactivates the DNase. Finally, multiple Csm3 subunits cleave the target transcripts, and release of target RNA cleavage products restores the binary conformation, completing the spatiotemporal regulation of LdCsm systems.

## Materials and Methods

### Bacterial strains and growth conditions

*L. delbrueckii* subsp. *bulgaricus* ND04 (GenBank: CP016393.1) was grown in MRS broth (Oxoid, UK) at 37°C without shaking. *E. coli* JM109 and BL21(DE3) were propagated in Luria-Bertani (LB) medium at 37°C with 200 rpm/min shaking. If applicable, antibiotics were added as the following: ampicillin (100 μg/ml, sigma), kanamycin (25 μg/ml, sigma) and chloramphenicol (10 μg/ml, sigma).

### Construction of different vectors and plasmids

To construct a p15A replicon-based expression vector, the origin fragment was obtained from pTRKH2^63^, ampicillin resistance gene (Amp) was derived from pUC19 (New England Biolabs) whereas gene expression cassette were amplified from and pET30a (Novagen) by PCR, using three sets of primers (p15A-F and-R, Amp-F and −R, as well as LacI-F and T7-R) listed in Supplementary Table S1. Ligation of these DNA fragments generated the expression vector p15AIE (Supplementary Fig. S2a). The same strategy was employed to construct pUCE in which the origin was amplified from pUC19, chloramphenicol resistance gene was obtained from pCI372 whereas the expression cassette, from pET30a, using the primer sets of pUC, Cm and T7, respectively (Supplementary Table S1). Ligation of these DNA fragments gave pUCE (Supplementary Fig. S2b).

The vector for invader plasmid assay was constructed in two steps. First, fragment 1 containing the origin of pBR322 and kanamycin resistance gene (Kan) was obtained from pET30a by PCR (pBR-Kan primers); second, DNA fragments carrying an arabinose-inducible Bad promoter, multiple cloning sites (MCS) and transcriptional terminator (T) were generated by PCR from *E. coli* BL21(DE3) genome and plasmid pELX1^64^, using the primer sets of PBad and MCS-T listed in Supplementary Table S1, respectively, and they were then fused together by splicing overlapping extension PCR (SOE-PCR)^65^; third, ligation of the fragment 1 and the SOE DNA fragment yielded pBad vector. Then, three DNA fragments carrying *eGFP* gene were amplified from pEGFP-N1^66^ by PCR using GFR-F/GFR-R, CTR-GFR-F/GFR-R and NTR-GFR-F/GFR-R, and insertion of each fragment into pBad individually gave pBad-G, pBad-CTR and pPBad-NTR. All primers employed in this work were listed in Supplementary Table S1.

Chromosomal DNA was extracted from cells of *L. delbrueckii* subsp. *bulgaricus* ND04 using OMEGA Genomic DNA Purification Kit (OMEGA Bio-tek). DNA fragment covering the cas6-cas10-csm2-csm3-csm4-csm5 gene cassette was amplified by PCR with the primers *Sal*I-*Cas6*-F and *Not*I-*Csm5*-R using ND04 genome DNA as template. The PCR product inserted into p15AIE via *Sal*I and *Not*I, yielding p15AIE-Cas. *csm2* gene was amplified from ND04 genome using primers Csm2-F and Csm2-R. PCR product was then digested with *Nde*I and *Xho*I and cloned into pET30a expression vectors, giving pET30a-Csm2. To construct a CRISPR array plasmid, fusion PCR amplification was performed using three primers Re-S1-F, S1-R1, Re-S1-R to generate the multiple copies of 36 nt length repeats interspaced by multiple S1 spacer (40 nt) of identical sequence, then PCR products of ~1 kb were recovered from an agarose gel using an OMEGA gel-purification kit (OMEGA Bio-tek). The purified DNA fragments were cloned into pJET1.2 (CloneJET PCR Cloning Kit, Thermo Scientific). After confirming the sequence of the synthetic CRISPR array on pJET clones (GATC Bio-tech), the DNA fragment was amplified and cloned into plasmid pUCE at the *Bgl*II site, yielding pUCE-S1 carried 10 identical spacers (S1) in the CRISPR array. Finally, plasmid p15AIE-Cas-S1 was constructed by insertion of the CRISPR array into p15AIE-Cas at the *Nhe*I site.

### Purification of LdCsm effector complexes from *E. coli*

Three plasmids (p15AIE-Cas, pUCE-S1 and pET30a-Csm2) were introduced into *E. coli* BL21 (DE3) by electroporation, yielding a *E. coli* strain containing all three plasmids. This bacterial strain was employed as host to overexpress the LdCsm system. The strain was cultured 200 ml LB medium containing ampicillin, kanamycin, chloramphenicol (at 37°C, 200 rpm) to the mid-log phase (OD_600_ was 0.8), then IPTG was added to 0.3 mM and the culture was further cultured at 25°C for 16 h. Cells were harvested by centrifugation at 5,000 rpm for 5 min, and cell pellets were resuspended in 20 ml buffer A [20 mM Tris-HCl (pH 8.5), 0.25 M NaCl, 20 mM imidazole and 10% glycerol], yielding cell suspension that was treated with a French press for cell lysis at 4°C. Cell debris was then removed from treated cell suspension by centrifugation at 10,000 rpm for 30 min at 4°C. The Csm complex was captured on the HiTrap affinity column (GE Healthcare) by LdCsm2 copurification and eluted with buffer B [20 mM Tris-HCl (pH 8.5), 0.25 M NaCl, 200 mM imidazole and 10% glycerol]. The resulting LdCsm effector complex preparation was further purified by size exclusion chromatography (SEC) with Superdex 200 (GE Healthcare) using the chromatography buffer [20 mM Tris-HCl (pH 8.5), 0.25 M NaCl and 5% glycerol]. SEC fraction samples were analyzed by sodium dodecyl sulfate-polyacrylamide gel electrophoresis (SDS-PAGE) and those containing the complete set and high quality of Csm complex components were pooled together and used for further analysis. Csm complex concentration was measured according to the Bradford method using the protein assay kit (Thermo Scientific) with bovine serum albumin as the standard.

### Extraction and analysis of crRNA

The purified LdCsm complex (100 μl) was first mixed with l00 μl Trizol agent (Sigma), and then 200 μl chloroform:isoamylalcohol (24:1, v/v) was added. After vortex for 30 s, the mixture was centrifuged at 12,000 rpm for 10 min at 4 °C. The upper phase was transferred into a new tube and reextracted with 200 μl chloroform:isoamylalcohol. crRNA in the upper phase was precipitated with one volume of isopropanol and washed twice with 1 ml of 70% ice-cold ethanol. The pellet was air-dried for 30 min at the room temperature and dissolved in 20 μl DEPC-H_2_O. Ten nanograms of crRNA was 5′-labeled with [γ-^32^P]-ATP (PerkinElmer) using T4 polynucleotide kinase (New England Biolabs) and separated on a 12% denaturing polyacrylamide gel. The labeled crRNAs were identified by exposing the gel to a phosphor screen (GE Healthcare) and scanned with a Typhoon FLA 7000 (GE Healthcare). For northern blotting of crRNA, 100 ng of unlabeled crRNA was mixed with equal volume of 2 × RNA loading dye (New England Biolabs) and fractionated in the 12% denaturing polyacrylamide gel. Northern blotting analysis was conducted as described previously^67^, using radiolabeled RNA S1-40 (Supplementary Table S1).

### Labeling of DNA and RNA Substrates

All DNA, S10 (nonhomologous RNA), S1-40 (PTR), S1-46 (CTR) and S1-48 (NTR) oligonucleotides were purchased from IDT, other RNA oligonucleotides were generated by *in vitro* transcription using TranscriptAid T7 High Yield Transcription Kit (Thermo Scientific) (Supplementary Table S1). DNA and RNA oligonucleotides to be used as substrate for cleavage and binding assays were purified by recovering the corresponding bands from either a native polyacrylamide gel (for double-stranded DNA) or from denaturing polyacrylamide gel (for ssDNA or RNA) after electrophoresis. ssDNA and RNA substrates were 5′ labeled with [γ-^32^P]-ATP and T4 polynucleotide kinase followed by denaturing gel purification. Double strand DNA, bubble DNA and R-loop DNA was generated as described previously^31^.

### Cleavage assay

Nucleic acid cleavage assays were conducted in 10 μl of reaction containing the indicated amount of effector complex and substrates in the cleavage buffer (50 mM Tris-Cl (pH 6.8), 10 mM MgCl_2_, 50 mM KCl, 0.1 mg/ml BSA]. In DNA cleavage assay, 500 nM (unless otherwise indicated) unlabeled RNA was supplemented to activate DNA cleavage activity. Samples were incubated at 37°C and stopped for indicated time periods and the reaction was stopped by addition of 2 × RNA loading dye (New England Biolabs). For electrophoresis, samples were heated for 3 min at 95°C and analyzed on an 18% polyacrylamide denaturing gel. RNA ladders were generated by Decade™ Marker RNA (Ambion) following the instructions and labeled by [γ-^32^P]-ATP with T4 polynucleotide kinase. Results were recorded by phosphor imaging.

### Determination of cyclic oligoadenylates synthesis activity

Each reaction mixture contained 50 nM Csm complex, 500 nM unlabeled S1-46 RNA (CTR), ~2 nM [α-^32^P]-ATP (PerkinElmer) and an indicated content of ATP in 50 mM Tris-Cl (pH 6.8) and 0.1 mg/ml BSA supplementing with ions. Reactions were incubated at 37°C for 120 min and 2 × RNA loading dye was added at the indicated time points to stop the reaction. Samples were kept on ice until use. For electrophoresis, samples were treated at 95°C for 3 min and analyzed by 24% denaturing PAGE. Gels were analyzed by phosphor imaging.

### Electrophoretic Mobility Shift Assay

ssDNA binding assay was performed by incubating different amounts of Csm complex (specified in each experiment) with 5 nM ^32^P-5′-labeled S10-60 ssDNA in the cleavage buffer. All reactions were incubated at 37°C for the indicated time periods. Then, the same volume of 2 × native loading buffer [0.1% bromophenol blue, 15% sucrose, w/v] was added and the samples were immediately put on ice and kept there until needed for electrophoresis on an 8% nondenaturing polyacrylamide gel. Electrophoresis was carried out at 4°C using 40 mM Tris, 20 mM acetic acid (pH 8.4 at 25°C) as the running buffer. Gels were analyzed by phosphor imaging.

Relative ssDNA binding and cleavage activities of LdCsm facilitated by each of these target RNAs were estimated by image quantification of bands on non-denaturing PAGE and denaturing PAGE, respectively, using the accessory analysis tool equipped with a Typhoon FLA 7000. Results of average of three independent assays are shown with bars representing the mean standard deviation (± SD).

### Mutagenesis of LdCsm1 and LdCsm3

*LdCsm1* mutants were generated using the splicing overlapping extension PCR protocol previously reported ^63^. In brief, several mutations were designed in the internal partial overlapping primers, initial PCRs were preformed using the external primer and their corresponding internal primers to generate overlapping gene segments, then, the two PCR products were fused together by overlapping extension PCR. The resulting fragments were digested with restriction endonuclease (DNA fragment containing LdCsm1 H15D16A mutation was cleaved with *Sal*I and *Sac*I, those carrying LdCsm1 E415A, E415C416A, C416C419A, D541D543A, D599D600A or Q597G were digested with *Stu*I and *Sac*I. LdCsm3 H20A, D34A and D106A was obtained via *Stu*I and *Kpn*I. After purification, these DNA fragments were inserted into the plasmid p15AIE-Cas at the corresponding restriction sites, yielding the plasmids carrying each designed *LdCsm1* mutation. All mutations were verified by DNA sequencing (GATC Biotech).

### Plasmid interference assay

Plasmid interference assays were performed as previously described^51^. Briefly, an *E. coli* BL21(DE3) strain carrying p15AIE-Cas-S1 (80 μl) was transformed with 100 ng of one of the following plasmids, pBad-G, pBad-CTR or pBad-NTR. Electroporation was performed in a 1-mm cuvette (Bio-Rad, USA), with the setting of 1,600 V, 200 Ω and 25 μF, using a Gene Pulser II Electroporation System (Bio-Rad). Then, 920 μl of SOC medium was immediately added to electroporated cells and incubated with shaking (200 rpm) at 37°C for 60 min. A series of dilutions were then made for each transformation, and 100 μl of each dilution was plated onto LB agar plates containing 0.05 mM IPTG, Ampicillin, Kanamycin and various concentrations of L-arabinose. Plates were incubated overnight at 37°C and the transformation efficiency was calculated. Transformation experiments were conducted for three independent times.

### Fluorescence assay

The reaction mixture (20 μl in total) contains 50 nM complex, the indicated concentration target RNA, in the presence of 500 nM of FAM-poly-16T-BHQ1 ssDNA substrate (Tsingke biotechnology company, Wuhan, China). These reactions were incubated in a 384-well black plate (Thermo fisher) and put on a fluorescence plate reader (FLUOstar Omega) for up to 60 min at 37 °C with fluorescence measurements taken every 1 min (λ_ex_: 485 nm; λ_em_: 535 nm). Background fluorescence values were obtained by subtracting fluorescence values obtained from reactions carried out in the absence of target RNA.

## Acknowledgments

We thank colleagues in the Archaea Centre for stimulating discussions and Dr. Søren M. Madsen (Bioneer A/S, Hørsholm, Denmark) for a kind gift of pTRKH2 and pCI372 plasmids. This work has been supported by grants from Danish Council for Independent Research (DFF-4181-00274), National Science Foundation of China (31771380) and the National Transgenic Science and Technology Program (2019ZX08010-003) to Q.S.

## Author contributions

H.Z. provided the *L. delbrueckii* subsp. *bulgaricus* strain; J.L., and Q.S. designed experiments; J.L. and M.F. performed experiments; J.L., M.F. and Q.S. analyzed data; J.L. and Q.S. wrote the paper. All authors read and approved the final manuscript.

## Additional information

Supplemental Information includes Supplemental figures and Supplemental tables can be found with this article online.

