## Supplemental information for "Characterization of a novel type III CRISPR-Cas effector provides new insights into the allosteric activation and suppression of the Cas10 DNase"

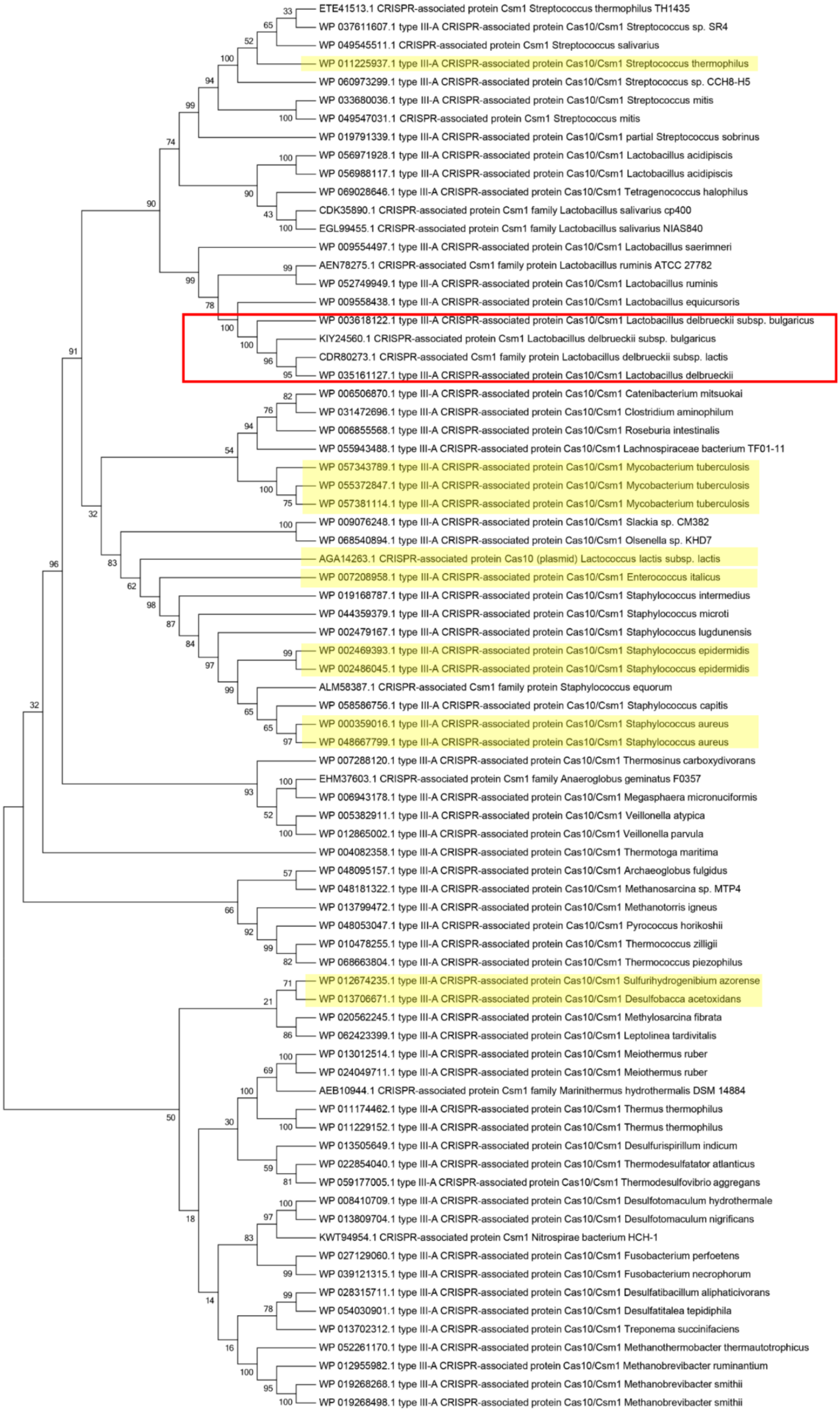


**Supplementary Fig. S1** The phylogenetic tree of the type III-A CRISPR-Cas system signature protein Cas10/Csm1 using MUSCLE method in MEGA5[1]. *L. delbrueckii* subsp. was highlighted by the red line box. The characterized type III-A system were highlighted by yellow background.

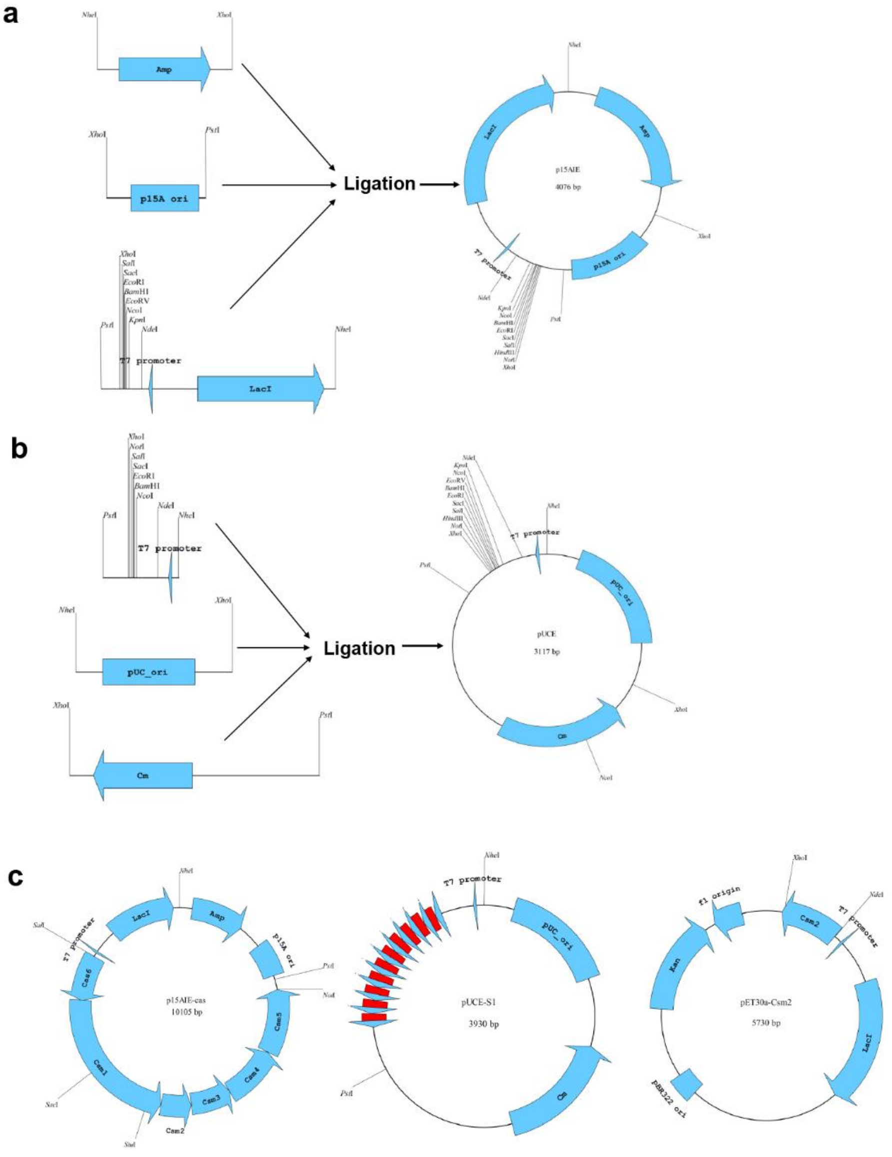


**Supplementary Fig. S2** Construction of expression vectors and plasmids.

(**a**) Construction of CRISPR-Cas cassette gene expression vector p15AIE, the replicon of p15A, ampicillin resistance gene (*Amp*) and gene expression/regulation elements (including T7 promoter, multiple cloning sites, T7 transcriptional terminator and regulation gene *LacI*) were amplified from plasmids pTRKH2, pUC19 and pET30a, respectively. Then, these three PCR products were ligated together in a triple ligation, yielding the expression vector p15AIE.

(**b**) Construction of crRNA expression vector pUCE. The replicon of pUC, chloramphenicol resistance gene (Cm) and gene expression elements (including T7 promoter, multiple cloning sites, T7 transcriptional terminator) were amplified from plasmids pUC19, pCI372, and pET30a, respectively. Then, these three DNA fragments were ligated together in a triple ligation, resulting in the expression vector pUCE.

(**c**) DNA fragment covering the *cas6-cas10-csm2-csm3-csm4-csm5* gene cassette was amplified from genome of *L. delbrueckii* subsp. *bulgaricus* ND04. The PCR product was ligated to plasmid vector p15AIE, yielding p15AIE-cas. For construction of plasmid pUCE-S1, fusion PCR amplification was performed to generate the multiply 36 nt length repeats interspaced by multiply identical 40 nt spacer S1, then the PCR products of ~1 kb were recovered by agarose electrophoresis and the OMEGA gel-purification kit. The obtained DNA fragments were cloned into a blunt ligation vector pJET1.2/blunt and confirmed by DNA sequencing, then the target DNA fragment was amplified and cloned into plasmid pUCE to generate the plasmid pUCE-S1 carried ten identical tandem copies of spacer S1. Finally, *csm2* gene was amplified from strain ND04 genome, the yielding PCR products were cloned into pET30a expression vectors to generate pET30a-Csm2.

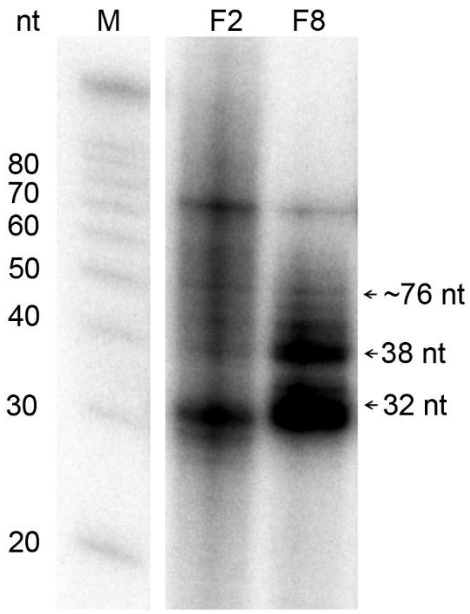


**Supplementary Fig. S3** Northern blot analysis of crRNA within the LdCsm effector complexes. RNAs were extracted from the SEC-purified LdCsm samples. M: RNA size ladder.

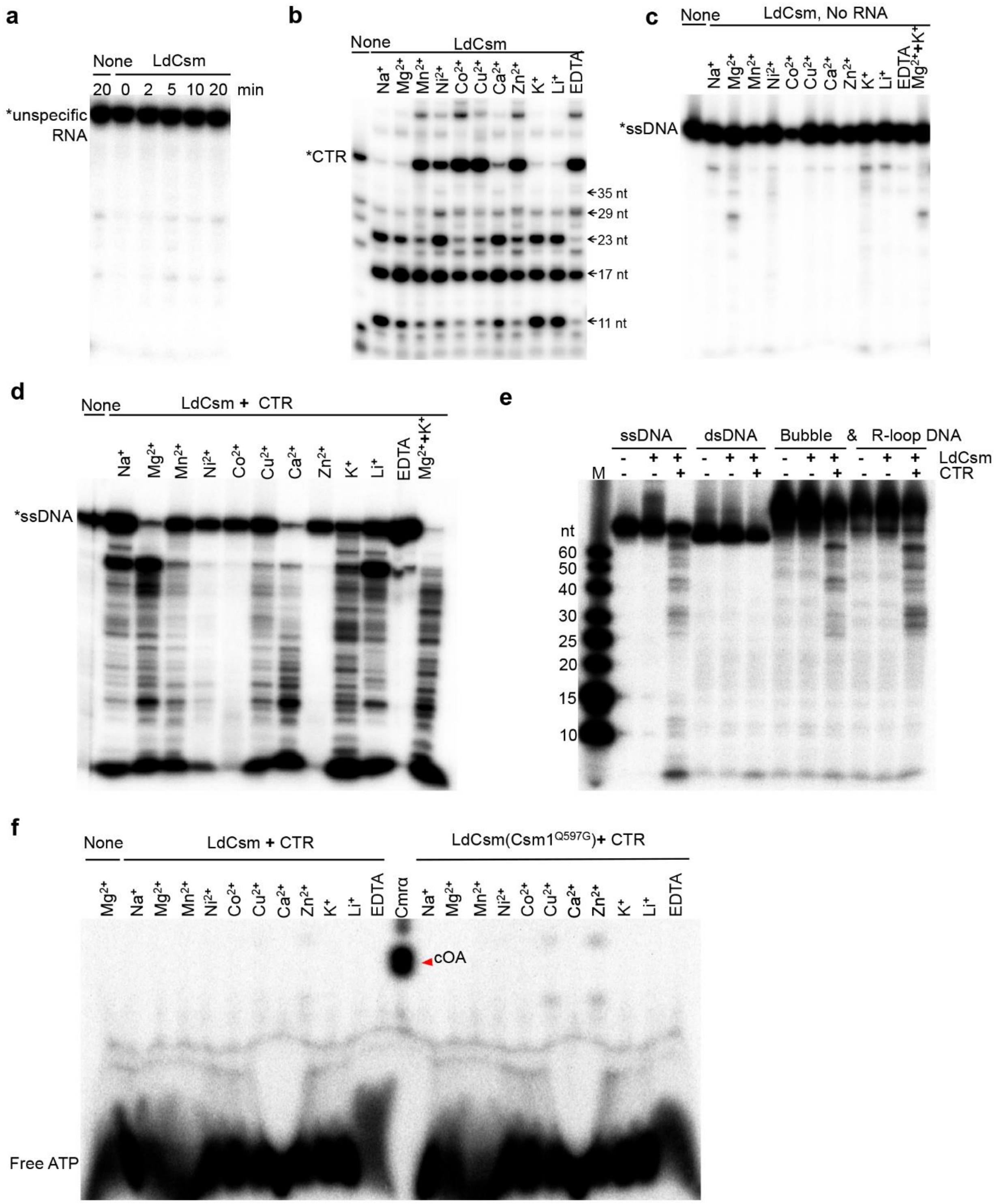


**Supplementary Fig. S4** Biochemical characterization of the LdCsm effector complexes**.**

(**a**) No non-homologous RNA cleavage by LdCsm. 50 nM of S10, a non-homologous RNA, was incubated with or without 50 nM of the LdCsm for the indicated time, then, the samples were analyzed by denaturing PAGE.

(**b**) Target RNA cleavage required cations. 50 nM of S1-46 target RNA (CTR) substrates were incubated with 50 nM of LdCsm addition with different ions for 10 min, then, the samples were analyzed by denaturing PAGE. The final concentrations of ions ($Mg^{2+}$, $Mn^{2+}$, $Ni^{2+}$, $Cu^{2+}$, $Co^{2+}$, $Ca^{2+}$ and $Zn^{2+}$) were 10 mM, the final concentrations of $Na^{+}$, $K^{+}$, $Li^{+}$ and EDTA were 50 mM, all the samples also contain an extra 25 mM Na+ from storage buffer for storage of LdCsm complex.

(**c**) No ssDNase activity without target RNA. 50 nM of S10-60 ssDNA substrates and 50 nM of LdCsm were incubated without any RNA in different ions for 10 min, then, the samples were analyzed by denaturing PAGE. The concentrations of ions used were identical to that used in (b).

(**d**) Target RNA-activated ssDNA cleavage required cations. 50 nM of S10-60 ssDNA substrates and 50 nM of LdCsm were incubated with 500 nM of CTR in different ions for 10 min, then, the samples were analyzed by denaturing PAGE. The concentrations of ions used were identical to that used in (b).

(**e**) LdCsm cleaved ssDNA and ssDNA region of bubble DNA and R-loop DNA. ssDNA, dsDNA, bubble DNA and R-loop DNA together with (+) or without (-) 10 nM of the LdCsm in the presence (+) or absence (-) of 500 nM of CTR were incubated for 10 min, the samples were analyzed by denaturing PAGE.

(**f**) No cOA generated by LdCsm. 2 nM α32P-ATP and 100 nM cold ATP were incubated with 50 nM LdCsm or LdCsm(Csm1$^{Q597G}$)complex in the presence of 500 nM CTR for 120 min in different ions, then the samples were analyzed by denaturing PAGE. The concentrations of ions used were identical to that used in (b).

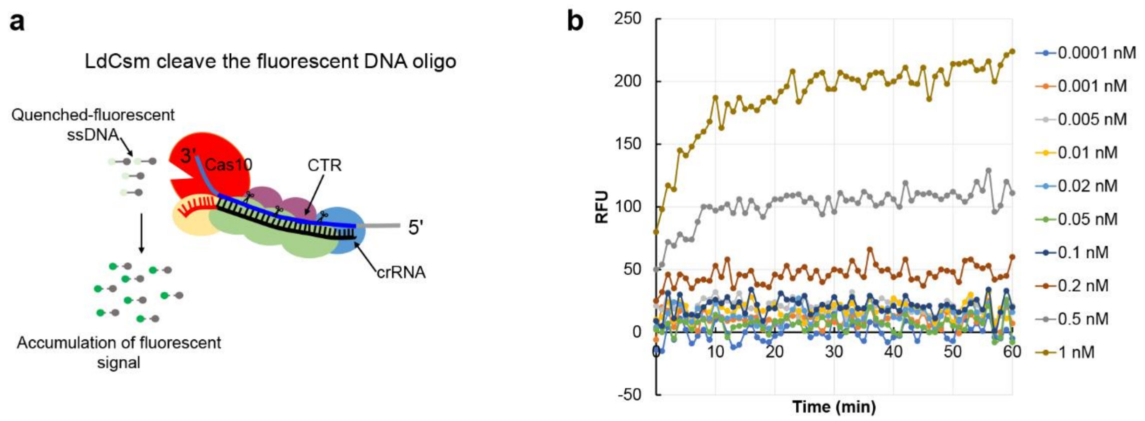


**Supplementary Fig. S5** The minimal concentration of target RNA requirement for the ssDNase activity of LdCsm. (**a**) Schematic of LdCsm RNA detection approach using a quenched fluorescent ssDNA oligo. (**b**) Quantified time-course fluorescence signal generated by LdCsm with varying concentrations of S1-46 RNA (CTR). Reactions (in total 20 μl) contain 500 nM of the FAM-poly-16T-BHQ1, varying concentrations of CTR, 50 nM of Csm effector.

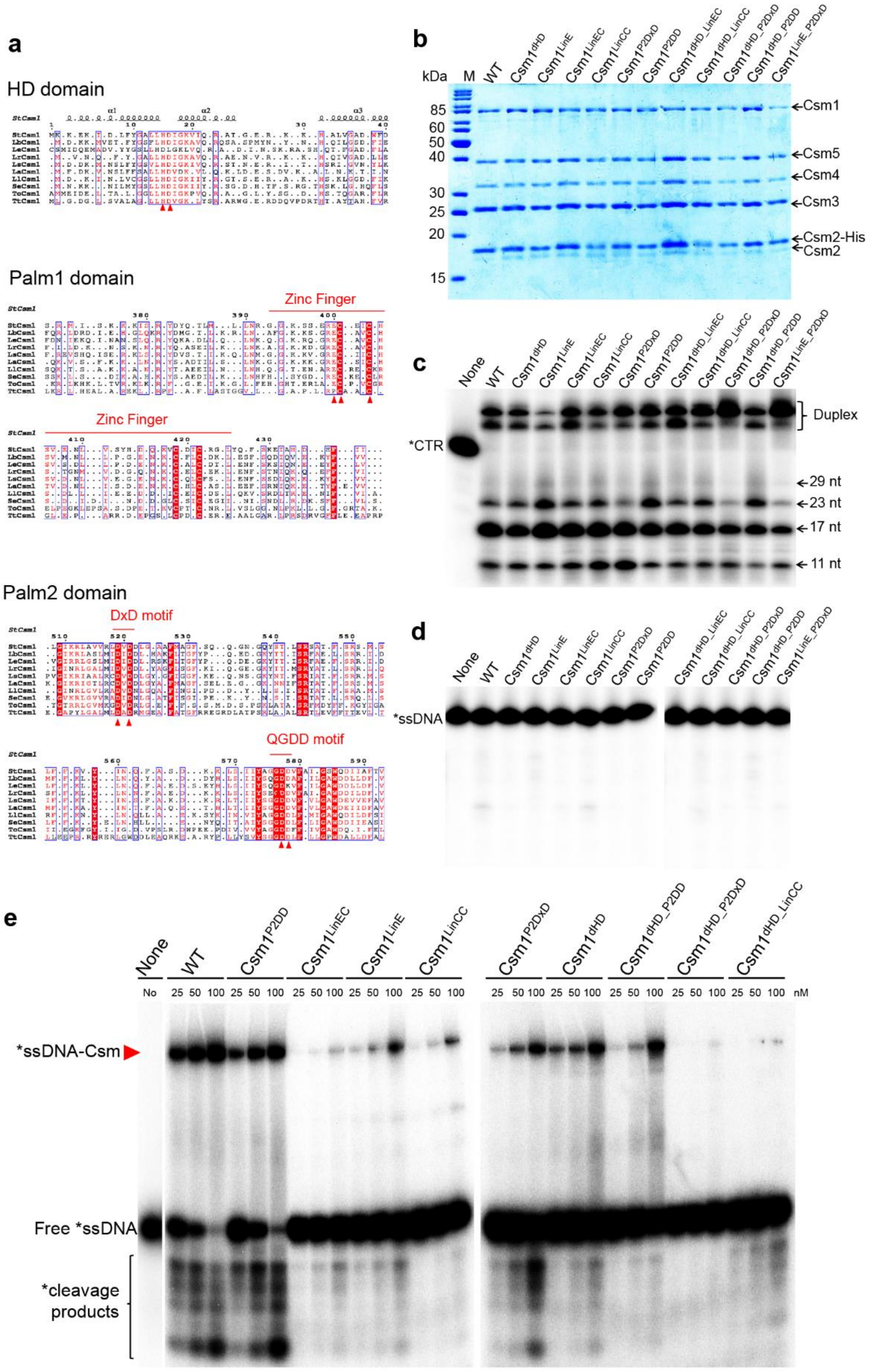


**Supplementary Fig. S6** Effect of LdCsm1 mutations on the target RNA cleavage, ssDNA cleavage and binding of LdCsm.

(**a**) Conserved residues in LdCsm1. Ten Csm1 homologues: *Lactobacillus bulgaricus* (Lb), *Streptococcus* *thermophilus* (St), *Lactobacillus equicursoris* (Le), *Lactobacillus ruminis* (Lr), *Lactobacillus salivarius* (Ls), *Lactobacillus acidipiscis* (La), *Lactococcus lactis* subsp. *lactis* (Ll), *Staphylococcus epidermidis* (Se), *Thermococcus onnurineus* (To) and *Thermus thermophilus* (Tt) were selected and aligned using MEGA5[1] and visualized using ESPript 3[2]. Some highly conserved residues marked with red arrows were chosen to produce the LdCsm1 variants.

(**b**) Coomassie blue-stained SDS-PAGE of protein components of the LdCsm1 mutated complexes, M: protein mass marker.

(**c**) Target RNA cleavage of LdCsm1 mutated variants. 50 nM of S1-46 target RNA (CTR) substrates were incubated with 50 nM of the indicated Csm complexes for 10 min, then, the samples were analyzed by denaturing PAGE. Duplex: Duplex of crRNA and substrate.

(**d**) No unspecific ssDNA cleavage without target RNA. 50 nM of S10-60 ssDNA substrates were incubated with 50 nM the indicated LdCsm1 mutated variants in the absence of target RNA for 10 min and the samples were analyzed by denaturing PAGE.

(**e**) Effect of LdCsm1 mutations on the ssDNA binding of LdCsm. 5 nM of S10-60 ssDNA substrates were incubated with the indicated concentration Csm in the presence of 500 nM of CTR for 3 min and then the samples were analyzed by non-denaturing PAGE, Red arrows represent the Csm-ssDNA binding complexes.

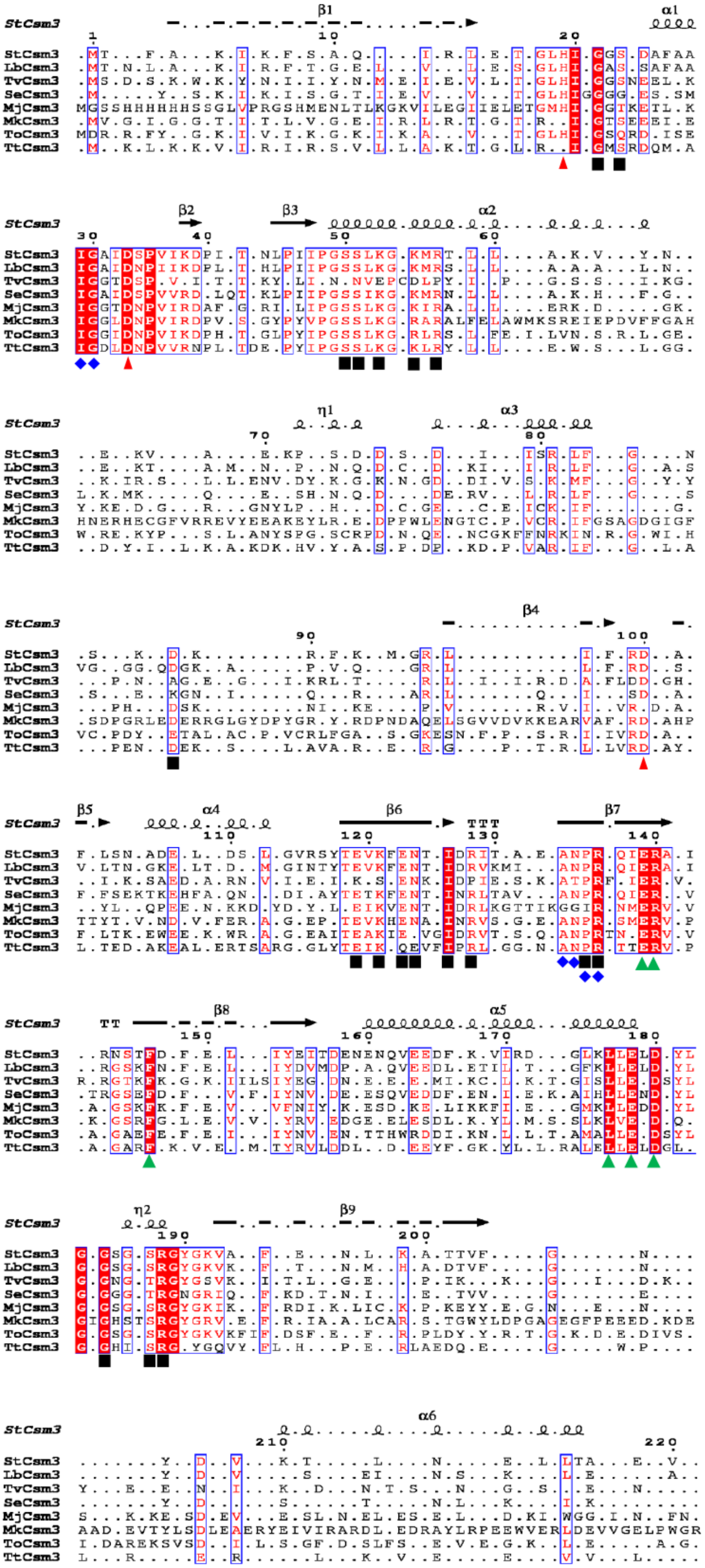
StCsm3
β1
α1
β2
β3
α2
η1
α3
β4
β5
α4
β6
β7
β8
α5
η2
β9
α6
StCsm3
LbCsm3
TvCsm3
SeCsm3
MjCsm3
MkCsm3
ToCsm3
TtCsm3

**Supplementary Fig. S7** Conserved residues in LdCsm3. Eight Csm3 homologues: *Lactobacillus bulgaricus* (Lb), *Streptococcus thermophilus* (St), *Thermoplasma volcanium* (Tv), *Staphylococcus epidermidis* (Se), *Methanocaldococcus jannaschii* (Mj), *Methanopyrus kandleri* (Mk), *Thermococcus onnurineus* (To) and *Thermus thermophilus* (Tt) were selected and aligned using MEGA5[1] and visualized using ESPript 3[2]. Some highly conserved residues were chosen to produce 5 LdCsm3 variants as indicated below: H20A ($Csm3^{H20A}$), D34A ($Csm3^{D34A}$), D106A ($Csm3^{D106A}$), the double substitutions H20/D34A ($Csm3^{H20/D34A}$) and D34/D106A ($Csm3^{D34/D106-A}$). The amino acid residues marked with red arrows were selected to construct these variants. The amino acid residues involve crRNA binding, target RNA binding and structural function were marked with black squares, blue rhombuses and green triangles, respectively.

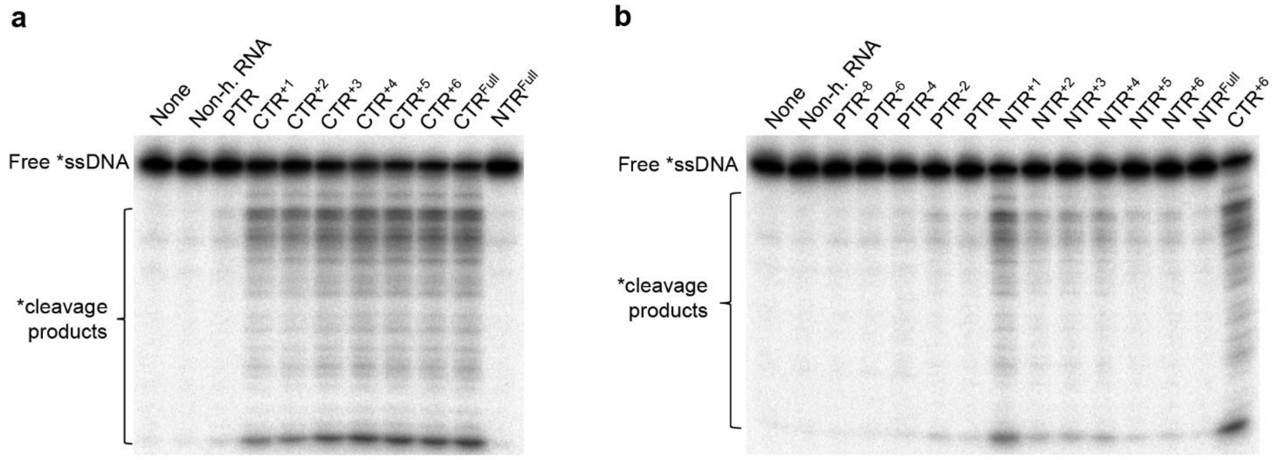


**Supplementary Fig. S8** DNase activity of LdCsm by target RNA involves activation and deactivation.

(**a**) CTR activates the LdCsm DNase. (**b**) NTR mediates autoimmunity avoidance by deactivation. Reactions were set up with 5 nM S10-60 ssDNA, 100 nM of LdCsm and 500 nM target RNA. After addition of one of target RNAs, the mixture was incubated at 37 °C for 3 min. Samples were then analyzed by denaturing PAGE. The percentage ssDNA cleaved of LdCsm facilitated by each of these target RNAs were estimated by image quantification of bands and showed in Fig. 5, using the accessory analysis tool equipped with a Typhoon FLA 7000. Results of average of three independent assays are shown with bars representing the mean standard deviation (± SD).

**Supplementary Table S1** Nucleic acid oligos and substrates used in this study.

| ssDNA and RNA | Sequence (5'-3') | Description |
|---|---|---|
| **oligo** | | |
| p15A-F | AGCCTGCAGGCGGTTTGCGTATTGGCTAG | For plasmid p15AIE construction |
| p15A-R | GGCCCTCGAGAAAAAAGGCTGCACCGG | For plasmid p15AIE construction |
| AMP-F | CTAGGCTAGCGACGAAAGGGCCTCGTGATAC | For plasmid p15AIE construction |
| Amp-R | GGCCCTCGAGCAGCAGATTACGCGCAG | For plasmid p15AIE construction |
| LacI-F | CTAGGCTAGCCACCGGAAGGAGCTGACTG | For plasmid p15AIE construction |
| T7-F | CTAGGCTAGCTCCGGCGTAGAGGATCGAG | For plasmid pUCE construction |
| T7-R | AGCCTGCAGATCCGGATATAGTTCCTCC | For plasmids p15AIE and pUCE construction |
| Cm-F | AGCCTGCAGCTTAAGGAACGTACAGACGG | For plasmid pUCE construction |
| Cm-R | CGCCTCGAGCATGTGAGGCATTTTCGCTC | For plasmid pUCE construction |
| pUC_ori-F | CTAGGCTAGCCTGTCAGACCAAGTTTACTC | For plasmid pUCE construction |
| pUC_ori-R | GGCCCTCGAGCATTAATGAATCGGCCAACGC | For plasmid pUCE construction |
| SalI-Cas6-F | AGGGTCGACGTGCAGAAGATAAGTTTAGTTTG | For plasmid p15AIE-Cas construction |
| NotI-Csm5-R | ATGGGCGGCCGCTCATTTCTTTTCTTTGATGGT G | For plasmid p15AIE-Cas construction |
| Csm2-F | GAATTCCATATGTACGGAAGTCCACGTCC | For plasmid pET30a-Csm2 construction |
| Csm2-R | GGCCCTCGAGATCCTTTCCGCCGTAATACT | For plasmid pET30a-Csm2 construction |
| Re-S1-F | GCTAACAACTTATCTCCGCTAGAAGGAGACGAG AACTTCAAAGCTTA | For plasmid pUCE-S1 construction |
| Re-S1-R | GTTCTCGTCTCCTTCTAGCGGAGATAAGTTGTTA GCTGTTAAGTCTGGT | For plasmid pUCE-S1 construction |
| S1-R1 | AGCTGTTAAGTCTGGTTTCCCTCCAGGGTATCT AAGCTTTGAAGTT | For plasmid pUCE-S1 construction |
| pBR322&Kan-F | CCGGAATTCGTTTACAATTTCAGGTGGC | For plasmid pBad construction |
| pBR322&Kan-R | CCGGGAGCTCTGAGAGCCTTCAACCCAGTC | For plasmid pBad construction |
| PBad-F | CCGGGAGCTCAGAAACCAATTGTCCATATTG | For plasmid pBad construction |
| PBad-R | CGTTTCACTCCATCCAAAAAAACG | For plasmid pBad construction |
| MCS-T-F | TTTTGGATGGAGTGAAACGATGCACCATCATCAT CATCAT | For plasmid pBad construction |
| MCS-T-R | CCGGAATTCCAACAGAACCGTTTCTACTC | For plasmid pBad construction |
| eGFP-F | CCGCCCATGGCAAGGGCGAGGAGCTG | For plasmid pBad-G construction |
| eGFP-R | CGGGGTACCAATCTCGATGTTGTGGCGGATC | For plasmids pBad-G and pBad-CTR or pBad-NTR construction |
| S1-F | CCGCCCATGGAAGTCTGGTTTCCCTCCAGGGTA | For plasmids pBad-CTR and |

| | | |
|---|---|---|
| | TCTAAGCTTTG | pBad-NTR construction |
| CTR-GFP-F | CCCTCCAGGGTATCTAAGCTTTGAAAAAAAAAGCAAGGGCGAGGAGCTG | For plasmid pBad-CTR construction |
| NTR-GFP-F | CTCCAGGGTATCTAAGCTTTGAAGTTCTCGTCAGCAAGGGCGAGGAGCTG | For plasmid pBad-NTR construction |
| Csm1-H15D16A-F | GTTTCTGGCTGCTATTGGTAAAGCCGTCCAAAG | For LdCsm1 H15D16-A mutation |
| Csm1-H15D16A-R | TACCAATAGCAGCCAGAAACGATCCGTAGAAGG | For LdCsm1 H15D16-A mutation |
| Csm1-D599D600A-F | CACAAGGTGCTGCTGCCTTTATTTTAGGTGCCTG | For LdCsm1 D599D600-A mutation |
| Csm1-D599D600A-R | TAAAGGCAGCAGCACCTTGTGAATAGATAATAC | For LdCsm1 D599D600-A mutation |
| Csm3-H20A-F | GCGGCCTGGCTATTGGCGCAAGTTCAGCTTTTG | For LdCsm3 H20A and H20/D34-A mutations |
| Csm3-H20A-R | GCGCCAATAGCCAGGCCGCTTTCCAGTACC | For LdCsm3 H20A and H20/D34-A mutations |
| Csm3-D34A-F | GGTGCGATCGCCAACCCGATAATTAAGGATCC | For LdCsm3 D34A mutation |
| Csm3-D34A-R | ATCGGGTTGGCGATCGCACCAATAGCCGCA | For LdCsm3 D34A mutation |
| Csm3-D106A-F | TATTTAGAGCCAGTGTTCTGACTAATGGCA | For LdCsm3 D106A and D34/D106-A mutations |
| Csm3-D106A-R | AGAACACTGGCTCTAAATAAAAGCCGCCCT | For LdCsm3 D106A and D34/D106-A mutations |
| SacI-Csm1-F | CTATGCTGAGCTCTTCCAGAG | For LdCsm1 Palm1 and Palm 2 mutations |
| SacI-Csm1-R | CTCTGGAAGAGCTCAGCATAG | For LdCsm1 H15D16-A mutation |
| StuI-Csm1-F | CGAGGTAAGGCCTTCATTTAC | For LdCsm3 D34A mutation |
| StuI-Csm1-R | GTAAATGAAGGCCTTACCTCG | For LdCsm1 Palm1 and Palm 2 mutations |
| KpnI-Csm4-R | GACAGTGGTACCGCGTATAAGC | For LdCsm3 D34A mutation |
| Csm1N-E415C416A-F | AAAAGAGTGGCCGGGCGGCTGCAGTTTGTCATAGCGTTATG | For LdCsm1 E415C416-A mutation |
| Csm1N-E415C416A-R | TATGACAAACTGCAGCCGCCCGGCCACTCTTTTTGCCAAAG | For LdCsm1 E415C416-A mutation |
| Csm1N-Q597G-F | GAGTATTATCTATTCAGGAGGTGATGATGCCTTTATTTTAGG | For LdCsm1 Q597G mutation |
| Csm1N-Q597G-R | ATAAAGGCATCATCACCTCCTGAATAGATAATACTCAGATG | For LdCsm1 Q597G mutation |
| Csm1N-E415A-F | AAAAGAGTGGCCGGGCGTGTGCAGTTTGTCATAGCGTTATG | For LdCsm1 E415A mutation |
| Csm1N-E415A-R | TATGACAAACTGCACACGCCCGGCCACTCTTTT | For LdCsm1 E415A mutation |

| | | |
|---|---|---|
| | TGCCAAAG | |
| Csm1N-C416/419A-F | GGAGGCTGCAGTTGCTCATAGCGTTATGAATCTCC | For LdCsm1 C416/419-A mutation |
| Csm1N-C416/419A-R | TATGAGCAACTGCAGCCTCCCGGCCACTCTTTTGC | For LdCsm1 C416/419-A mutation |
| Csm1N-D541/543A-F | ATGATCGCTATTGCTGACCTCCATGCCAAATTCT | For LdCsm1 D541/543-A mutation |
| Csm1N-D541/543A-R | GAGGTCAGCAATAGCGATCATCAAAGAAGCAAG | For LdCsm1 D541/543-A mutation |
| **Nucleic acid substrates** | | |
| S10 RNA | AUAGAAUGCCCCCAUUAUACAAUAUCUACGUUUUAGAUGAAAAAAA | Non-homologous RNA |
| S1-40 (PTR) | UGUUAAGUCUGGUUUCCCUCCAGGGUAUCUAAGCUUUGAA | Target RNA Lacking any 3' anti-tag |
| S1-46 (CTR) | UGUUAAGUCUGGUUUCCCUCCAGGGUAUCUAAGCUUUGAAAAAAAA | Target RNA with noncomplementary 3' anti-tag with crRNA |
| S1-48 (NTR) | UGUUAAGUCUGGUUUCCCUCCAGGGUAUCUAAGCUUUGAAGUUCUCGU | Target RNA with complementary 3' anti-tag with crRNA |
| S10-60 ssDNA | ACTATAGGGAGAATAGAATGCCCCCATTATACAATATCTACGTTTTAGATGACCCCCCCC | Non-homologous ssDNA substrate |
| $PTR^{-2}$ | GGUGUUAAGUCUGGUUUCCCUCCAGGGUAUCUAAGCUUUG | A PTR Lacking double nucleotides in 3'-end |
| $PTR^{-4}$ | GGUGUUAAGUCUGGUUUCCCUCCAGGGUAUCUAAGCUU | A PTR Lacking four nucleotides in 3'-end |
| $PTR^{-6}$ | GGUGUUAAGUCUGGUUUCCCUCCAGGGUAUCUAAGC | A PTR Lacking six nucleotides in 3'-end |
| $PTR^{-8}$ | GGUGUUAAGUCUGGUUUCCCUCCAGGGUAUCUAA | A PTR Lacking eight nucleotides in 3'-end |
| $CTR^{+1}$ | GGUGUUAAGUCUGGUUUCCCUCCAGGGUAUCUAAGCUUUGAAA | Cognate target RNA with single A in 3' anti-tag |
| $CTR^{+2}$ | GGUGUUAAGUCUGGUUUCCCUCCAGGGUAUCUAAGCUUUGAAAA | Cognate target RNA with double A in 3' anti-tag |
| $CTR^{+3}$ | GGUGUUAAGUCUGGUUUCCCUCCAGGGUAUCUAAGCUUUGAAAAA | Cognate target RNA with triple A in 3' anti-tag |
| $CTR^{+4}$ | GGUGUUAAGUCUGGUUUCCCUCCAGGGUAUCUAAGCUUUGAAAAAA | Cognate target RNA with four A in 3' anti-tag |
| $CTR^{+5}$ | GGUGUUAAGUCUGGUUUCCCUCCAGGGUAUCUAAGCUUUGAAAAAAA | Cognate target RNA with five A in 3' anti-tag |
| $CTR^{+6}$ | GGUGUUAAGUCUGGUUUCCCUCCAGGGUAUCUAAGCUUUGAAAAAAAA | Cognate target RNA with six A in 3' anti-tag |

| | | |
|---|---|---|
| $CTR^{Full}$ | GGUGUUAAGUCUGGUUUCCCUCCAGGGUAUC UAAGCUUUGAAAAAAAAAA | Cognate target RNA with eight A in 3' anti-tag |
| $NTR^{+1}$ | GGUGUUAAGUCUGGUUUCCCUCCAGGGUAUC UAAGCUUUGAAG | Non-Cognate target RNA with single nucleotide in 3' anti-tag |
| $NTR^{+2}$ | GGUGUUAAGUCUGGUUUCCCUCCAGGGUAUC UAAGCUUUGAAGU | Non-Cognate target RNA with double nucleotides in 3' anti-tag region |
| $NTR^{+3}$ | GGUGUUAAGUCUGGUUUCCCUCCAGGGUAUC UAAGCUUUGAAGUU | Non-Cognate target RNA with triple nucleotides in 3' anti-tag |
| $NTR^{+4}$ | GGUGUUAAGUCUGGUUUCCCUCCAGGGUAUC UAAGCUUUGAAGUUC | Non-Cognate target RNA with four nucleotides in 3' anti-tag |
| $NTR^{+5}$ | GGUGUUAAGUCUGGUUUCCCUCCAGGGUAUC UAAGCUUUGAAGUUCU | Non-Cognate target RNA with five nucleotides in 3' anti-tag |
| $NTR^{+6}$ | GGUGUUAAGUCUGGUUUCCCUCCAGGGUAUC UAAGCUUUGAAGUUCUC | Non-Cognate target RNA with six nucleotides in 3' anti-tag |
| $NTR^{Full}$ | GGUGUUAAGUCUGGUUUCCCUCCAGGGUAUC UAAGCUUUGAAGUUCUCGU | Non-Cognate target RNA with eight nucleotides in 3' anti-tag |

**Supplementary Table S2** LdCsm1 and LdCsm3 mutations utilized in this study.

| RNP Name | Description |
|---|---|
| $Csm1^{dHD}$ | A HD domain mutant carrying H15A and D16A substitutions |
| $Csm1^{LinE}$ | A Linker domain mutant carrying E415A substitution |
| $Csm1^{LinEC}$ | A Linker domain mutant carrying E415A and C416A substitution |
| $Csm1^{LinCC}$ | A Linker domain mutant carrying C416A and C419A substitutions |
| $Csm1^{P2DxD}$ | A Palm2 domain mutant carrying D541A and D543A substitutions |
| $Csm1^{P2DD}$ | A Palm2 domain mutant carrying D599A and D600A substitutions |
| $Csm1^{dHD\_LinEC}$ | A Csm1 mutant carrying H15A, D16A, E415A and C416A substitutions |
| $Csm1^{dHD\_LinCC}$ | A Csm1 mutant carrying H15A, D16A, C416A and C419A substitutions |
| $Csm1^{dHD\_P2DxD}$ | A Csm1 mutant carrying H15A, D16A, D541A and D543A substitutions |
| $Csm1^{dHD\_P2DD}$ | A Csm1 mutant carrying H15A, D16A, D599A and D600A substitutions |
| $Csm1^{LinE\_P2DxD}$ | A Csm1 mutant carrying E415A, D541A and D543A substitutions |
| $Csm1^{Q597G}$ | A Csm1 mutant carrying Q597G substitution |
| $Csm3^{H20A}$ | A Csm3 mutant carrying H20A substitution |

| | |
|---|---|
| Csm3$^{D34A}$ | A Csm3 mutant carrying D34A substitution |
| Csm3$^{D106A}$ | A Csm3 mutant carrying D106A substitution |
| Csm3$^{H20/D34-A}$ | A Csm3 mutant carrying H20A and D34A substitutions |
| Csm3$^{D34/106-A}$ | A Csm3 mutant carrying D34A and D106A substitutions |
